# Integrated transcriptome analysis of Huntington’s disease iPSC-derived and mouse astrocytes implicates dysregulated synaptogenesis, actin, and astrocyte maturation

**DOI:** 10.1101/2022.07.28.501170

**Authors:** Andrea M. Reyes-Ortiz, Edsel M. Abud, Mara S. Burns, Jie Wu, Sarah J. Hernandez, Nicolette Geller, Keona Q. Wang, Corey Schulz, Ricardo Miramontes, Alice Lau, Neethu Michael, Emily Miyoshi, Mathew Blurton-Jones, David Van Vactor, John C. Reidling, Vivek Swarup, Wayne W. Poon, Ryan G. Lim, Leslie M. Thompson

## Abstract

Huntington’s disease (HD) is a neurodegenerative disease caused by an expanded CAG repeat within the *Huntingtin* (*HTT*) gene having dysregulated cellular homeostasis in the central nervous system, particularly in the striatum and cortex. Astrocytes establish and maintain neuronal functions through the secretion of soluble factors and physical interactions with other neurovascular unit cell types. Under pathological conditions, astrocytes can become reactive, causing cell state transitions that affect brain function. To investigate transitions between cellular states in unaffected and HD astrocytes at high resolution, single-nuclei RNA-sequencing (snRNA-seq) was performed on human HD patient induced pluripotent stem cell (iPSC)-derived astrocytes and on striatal and cortical tissue from a rapidly progressing HD mouse model (R6/2). Analysis of HD human and mouse astrocytes revealed both models have alterations in morphology, glutamate uptake, and dysregulation of astrocyte identity and maturation, whereas dysregulated actin-mediated signaling was unique to human iPSC-derived astrocytes. Representative proteins showed altered levels by Western. In both species, HD transcriptional changes reveal potential astrocyte maturation deficits that were potentially driven by astrogliogenesis transcription factors, including ATF3 and NFIA. When perturbed in a drosophila model of HD, knockdown of NFIA in glia rescued the climbing deficit. These data further support the hypothesis that mutant HTT induces dysregulated astrocyte cell states resulting in dysfunctional astrocytic properties, suggests that some of these states are cell autonomous and maybe unique to human HD, and implicate ATF3 and maturation deficits in HD pathogenesis.

## Introduction

Huntington’s disease (HD) is a devastating, autosomal-dominant neurodegenerative disease characterized by movement abnormalities, psychiatric disturbances, and cognitive impairment (Ross & Tabrizi, 2011) with no disease-modifying treatmentClick or tap here to enter text.. The genetic cause of HD is a CAG repeat expansion in the first exon of the *huntingtin* (*HTT*) gene, which encodes an expansion of a polyglutamine (polyQ) repeat tract in the Huntingtin (HTT) protein (The Huntington’s Disease Collaborative Research Group, 1993). When the *HTT* CAG repeat is expanded to 40 CAGs or above, HD is fully penetrant (Nance & Myers, 2001; Zuccato et al., 2010). HTT is ubiquitously expressed in all cell types in the body and a broad range of cellular processes, predominantly in the brain, are impacted by chronic mutant HTT (mHTT) expression (Bates et al., 2015; Ross & Tabrizi, 2011). The mutation alters HTT structure and function, causing toxic gain of function and loss of normal HTT functions(Bates et al., 2015). While the most overt HD neuropathology involves striatal neuronal loss and cortical atrophy, modeling of human induced pluripotent stem cells (iPSCs) has been leveraged to identify HD cell-autonomous defects in non-neuronal cell types, such as endothelial cells that comprise the blood-brain barrier (Lim et al., 2017), oligodendrocytes (Osipovitch et al., 2019), and astrocytes (Benraiss et al., 2021; Garcia et al., 2019; Juopperi et al., 2012; Osipovitch et al., 2019), indicating that chronic expression of mHTT may induce intrinsic deficits within each cell type.

Astrocytes are the major glial cell type in the CNS, playing a critical role in regulating brain homeostasis. Astrocytes provide neurotrophic support through glutamate uptake and recycling, and contribute to synaptogenesis, neuronal maturation, and neuronal maintenance. Inter-astrocytic communication of calcium signaling is a function of astrocytes thought to help control blood flow for coupling to neuronal energy demand (Khakh & Sofroniew, 2015; Zhao et al., 2015). In addition to regulating a microenvironment that facilitates neuronal signaling, astrocytes maintain direct interactions with the brain endothelium to establish and maintain BBB and neuronal functions through soluble and physical communications (Abbott et al., 2006).

There has been growing literature characterizing the development and regulation of unique cellular states that define functional states of astrocytes. Moreover, impairments in these astrocyte states may generate toxic functions that contribute to neurological disorders; however, the extent and mechanisms driving these impairments are just emerging for HD. HD astrocyte dysfunction appears to contribute to neuronal dyshomeostasis (Jiang et al., 2016; Khakh & Sofroniew, 2014; Tong et al., 2014), including impaired glutamate signaling and K+ buffering in astrocytes from HD mice (Diaz-Castro et al., 2019; Lee et al., 2013; Milnerwood et al., 2010; Tong et al., 2014). Glutamate related gene expression changes and dysfunction have recently been demonstrated in human patient post-mortem tissue (Diaz-Castro et al., 2019) and human patient iPSC-derived cells (Garcia et al., 2019), respectively. HD iPSC-derived astrocytes carrying juvenile-onset range CAG repeat lengths (77 CAG and 109 CAG) provided less support for functional neuronal maturation and contributed to glutamate-induced toxicity in iPSC-derived striatal neurons (Garcia et al., 2019). Other HD iPSC-astrocyte based studies revealed an increased autophagic response (Juopperi et al., 2012) and increased evoked inflammatory responses (Hsiao et al., 2015). Finally, purification of striatal astrocytes revealed common changes to RNA expression from R6/2 and Q175 HD mouse models, including core genes in calcium-dependent processes, G protein-coupled receptors and glutamate receptor signaling (Diaz-Castro et al., 2019) or in pathways regulating cytoskeletal formation and cell branching (Benraiss et al., 2021). R6/2 uniquely showed changes to cholesterol regulation (Benraiss et al., 2021). However, studies have yet to examine the molecular heterogeneity at a single cell level to define distinct astrocyte states from striatum and cortex in HD mouse models and patient iPSCs as well as potential mechanisms controlling their dysregulation.

With the development of single-cell and single-nuclei RNA-sequencing (sc- or snRNA-seq), astrocyte cell state transitions can now be compared at high resolution across multiple species, brain regions and ages. A recent snRNA-seq study on HD astrocytes from post-mortem patient cortical tissue identified heterogeneous cell states in HD human astrocytes (Al-Dalahmah et al., 2020). Increased expression of metallothionein and heat shock genes and loss of astrocyte-specific gene expression was demonstrated. To further investigate HD human astrocyte cell states, human patient iPSC-derived astrocytes allow the ability to ascertain earlier, cell autonomous events at the single-cell level that may lead to later changes in post-mortem human tissue. Furthermore, iPSC-derived astrocytes allow for functional assessment, linking transcriptional states to astrocyte dysfunction.

The goals of the present study are to further characterize astrocyte cell states in unaffected and HD contexts, identify drivers of transcriptional dysregulation, and to compare these states across human iPSC and mouse model systems. We investigated which astrocyte states exist in HD and if their transcriptional signatures can help predict altered astrocyte functions. We used unbiased snRNA-seq clustering followed by pathway enrichment and systematically compared signaling states between human and mouse astrocytes. Using this approach, we found altered astrocyte cell states relating to glutamate signaling and maturation common to both HD iPSC-derived astrocytes (iAstros) and striatal and cortical R6/2 astrocytes compared to unaffected control astrocytes. We also identified disease cell states that predict activated extracellular matrix (ECM) and dysregulated actin cytoskeletal dynamics to be unique to human HD astrocytes, informing cell-autonomous dysregulation related to endogenous CAG repeat expansion in astrocytes. These transcriptional signatures were then used to identify regulatory genes that may be driving these alterations and could represent therapeutic targets. Overall, HD transcriptional changes reveal potential astrocyte maturation deficits and implicate astrogliogenesis transcription factors, including ATF3 and NFIA, play a regulatory role contributing to HD pathogenesis. Together, this data provides a resource of astrocyte cell states and transitions that may be useful for broadening understanding the diverse astrocytes in the brain and guide therapeutic intervention in HD and suggests that loss of identity and maturation impairments are driven by altered expression of specific astrogliogenesis transcription factors.

## Results

### HD iAstros Exhibit Altered Morphology and Astrocyte Marker Expression

To understand how CAG repeat expansion may influence astrocyte function in HD, we evaluated heterogenous astrocyte cell states through quantifiable gene expression analysis at the single-cell level. We first investigated whether altered cell state changes are reflected through cell-intrinsic mechanisms in astrocytes differentiated from human HD versus control iPSCs that express *mHTT* in an endogenous context. HD (46Q, 53Q) and control iPSCs (18Q and 33Q) (HD iPSC Consortium, 2013, 2017; Lim et al., 2017; Smith-Geater et al., 2020) were differentiated into astrocytes (iAstros). Briefly, iPSCs were subject to prolonged neural induction to generate gliogenic neural stem cells followed by exposure to astrogliogenic factors for 60 days to induce morphologically and functionally mature astrocytes (**Figure 1A & S1**). HD iAstros had fewer astrocytic processes and had significantly larger cell volumes (**Figure S2**), indicating a potential deficit in astrocyte maturation (Zhang et al., 2016). We observed a significant decrease in the percentage of HD iAstros compared to control astrocytes that express GLAST, a functional glutamate transporter that clears the excitatory neurotransmitter from the extracellular space at synapses and is expressed by mature human astrocytes (**Figure 1C & D**). To directly compare a purer population of HD to control mature astrocytes, GLAST-positive iAstros were isolated by FACS sorting and used for all subsequent studies (**Figure 1B-D**). GLAST-sorted HD and control cells express canonical astrocyte markers, like GFAP, and exhibit a classical star-like morphology (**Figure 1F**).

**Figure 1.**
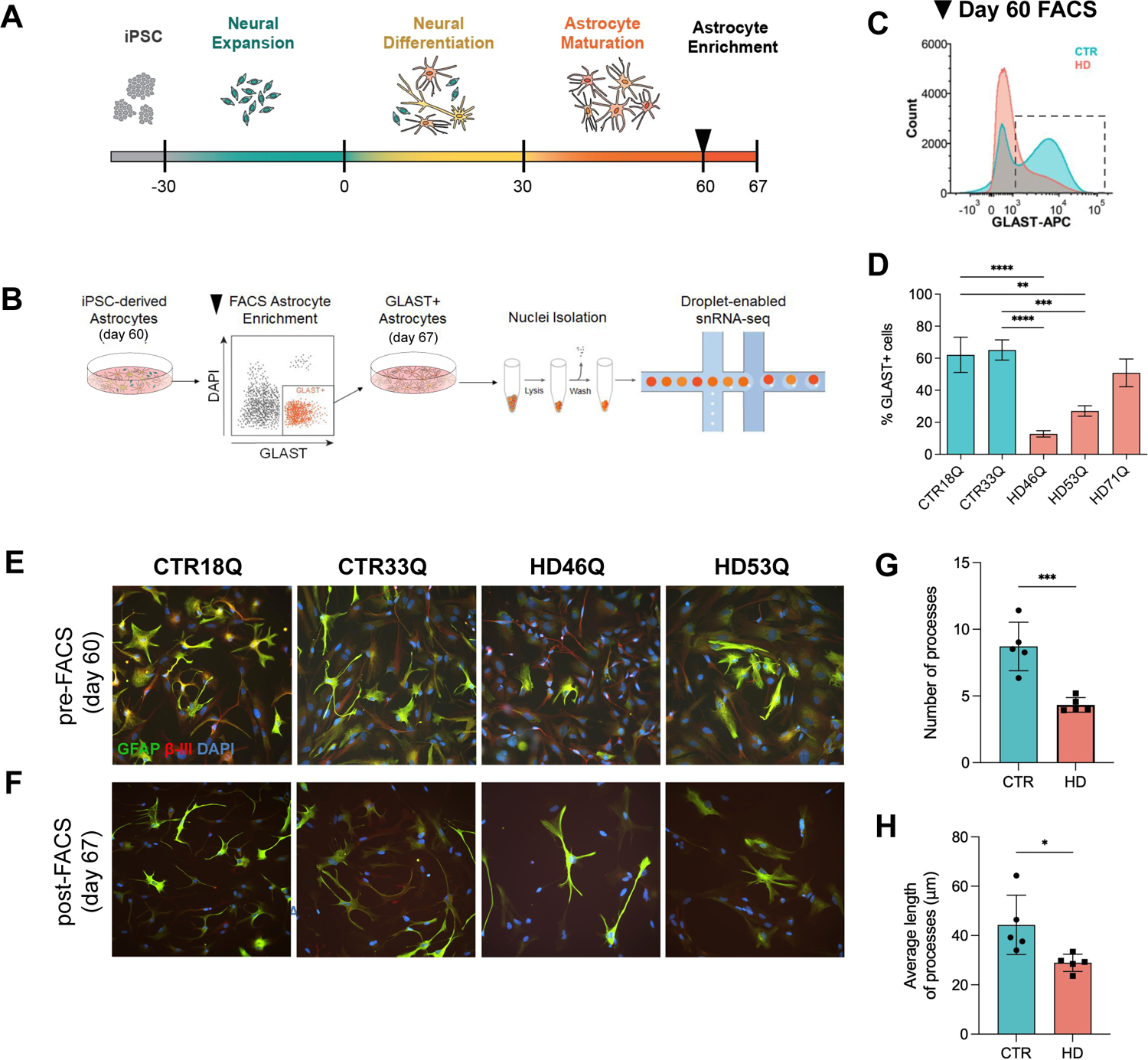
HD induced pluripotent stem cell-derived astrocyte derivation. (**A**) Astrocyte Differentiation first begins with the differentiation of iPSCs into gliogenic neural stem cells (NSCs), followed by 30 days of neural differentiation and 30 days of astrocyte maturation. (**B**) At day 60, GLAST+/DAPI-FACS was performed for astrocyte-enrichment and astrocytes are subsequently matured for 7 days. At day 67, GLAST+ astrocytes underwent single-nuclei isolation prior to droplet-enabled single-nuclei RNA-sequencing (snRNA-seq). (**C**) Representative FACS histogram of HD and control iAstros GLAST expression level demonstrates HD astrocytes have decreased expression of GLAST at day 60. (**D**) FACS quantification of GLAST+/DAPI-populations in day 60 iPSC-derived astrocytes (one-way ANOVA, 18Qv46Q ****p<0.0001 [n=8,11], 18Qv53Q **p=0.0028 [n=8,11], 33Qv46Q **p<0.0001 [n=8,11], 33Qv53Q ****p<0.0001 [n=8,11], 18Qv71Q ns [n=8,8], 33Qv71Q ns [n=8,8]). (**E-F**) Immunocytochemistry for GFAP, β-III tubulin (β-III), and DAPI at day 60 prior to FACS. (E) Immunocytochemistry for GFAP, SLC1A3, and DAPI 7 days post-FACS astrocyte enrichment (F). (**G-H**) HD iAstros have a significantly less total number of processes (unpaired t-test, CTRvHD ****p = 0.0009, n=2-3 biological [differentiation] replicate/line, n=5/genotype) (G) and significantly shorter average length of processes (unpaired t-test CTRvHD *p = 0.0253, n=2-3 biological [differentiation] replicate/line, n=5/genotype) (H). All error bars indicate ± standard error mean.

To further investigate morphological differences, we performed Sholl analysis on GLAST-sorted HD and control iAstros. HD GLAST-positive iAstros had less total number of astrocytic processes (**Figure 1G**) compared to GLAST-positive control iAstros. Additionally, HD GLAST-positive iAstros had shorter average length of processes (**Figure 1H**). Together, decreased length and number of astrocytic processes indicate that GLAST-positive HD astrocytes may have a decreased maturation profile compared to GLAST-positive control astrocytes.

### Astrocyte Transcriptional States in Patient-Derived iAstros Reveals Immature Cell States Regulated by Aberrant Astrogenesis Transcription in HD

Next, to investigate astrocyte transcriptional signatures in GLAST-sorted HD and control iAstros we used snRNA-seq (**Figure 1B**). Among the 36,128 nuclei sequenced, nine distinct cell clusters were identified across HD and control iAstros representing unique astrocyte cell states across Uniform Manifold Approximation and Projection (UMAP) plots (**Figure 2A**). HD or control-enriched clusters were identified by the number of iAstros per genotype by cluster (**Figure 2B**). Control-enriched cluster 0 highly expressed mature astrocyte markers (**Figure 2C**). Clusters 4 and 6 were also control-enriched (>60%). HD-enriched clusters 1, 2, 3, 5, and 8 expressed low levels of mature astrocyte markers, with cluster 3 expressing high levels of immature human astrocyte markers (*MKI67 and TOP2A*), as transcriptionally defined from fetal human astrocytes (Zhang et al., 2016). In addition to expression of immature astrocyte markers, HD-enriched cluster 8 had high expression of progenitor markers, like *PAX6*. Cluster 7 also highly expressed immature astrocyte markers but was not significantly enriched by genotype (∼59% HD). Based on gene expression profiling of astrocyte developmental markers, control and HD iAstros reflect most cell states including various developmental states, however, HD iAstros show clear shifts in composition and full depletion of one of the control states suggesting altered astrocyte maturation.

**Figure 2.**
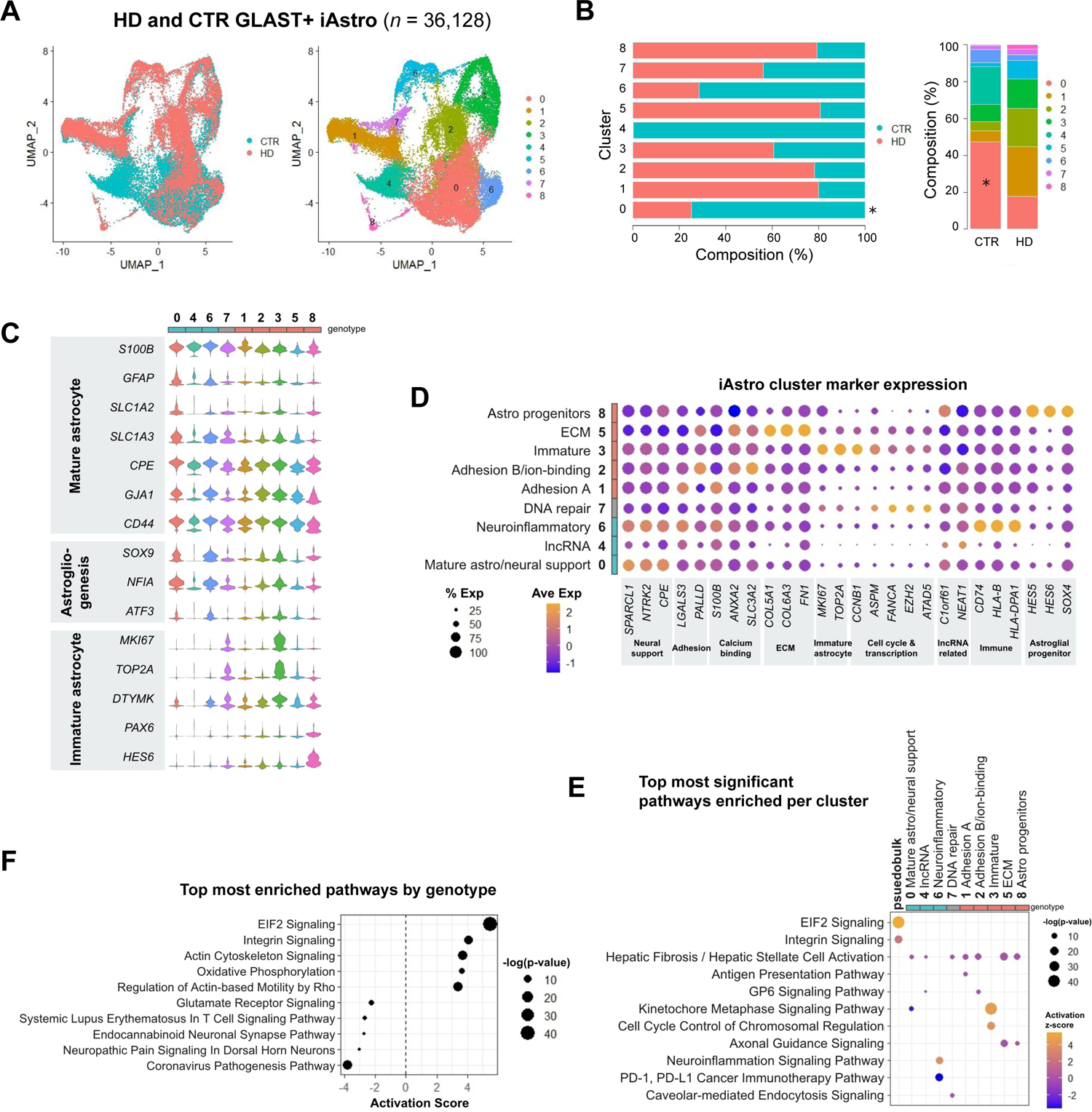
Dysregulated HD patient-derived astrocytes implicate immature cell states regulated by aberrant astrogliogenesis transcription. (**A**) GLAST+ iAstro snRNA-seq UMAP by genotype (CTR n=2, HD n=2) and by cluster. (**B**) Genotype composition of cells across astrocyte subclusters and by genotype (two-way ANOVA, Bonferroni multiple comparisons test performed on number of nuclei/mouse within each subcluster adjusted p-values: cluster 0 *p<0.05, cluster 1-8 ns; n=2 samples per genotype) (**C**) Astrocyte marker genes across each iAstro cluster shows a decrease in maturation gene expression in several HD-enriched subclusters. Genotype enrichment was defined by >60% of cluster, gray genotype represents 40-60%. (**D**) Top cluster markers for each cluster for cell state classification. (**E**) Top significant pathways across all iAstro clusters used to classify cell state signatures. Psuedobulk genotype pathway enrichment using DEGs from all HD iAstros compared to control iAstros is included for comparison. (**F**) Top 5 most significantly activated and inhibited pathways in HD iAstros.

To molecularly assess the unique transcriptional signatures across iAstro clusters, the top markers per cluster were plotted and classified into gene categories (**Figure 2D**). Control-enriched iAstro cluster 0 and 6 had the highest expression of neural support-related markers, like astrocyte-secreted synaptogenesis regulator *SPARCL1* and brain-derived neurotrophic factor (BDNF) receptor *NTRK2*. Interestingly, immune-related genes, including *CD74* and many HLAs, were also highly expressed in control-enriched iAstro cluster 6. Control-enriched cluster 4 represents a unique cell state with the most highly expressed genes related to long non-coding RNAs like *NEAT1,* and reduced expression of ECM and progenitor markers. HD-enriched clusters 1 and 2 had high expression of adhesion-related molecules, *LGALS3* and *PALLD*, with cluster 2 also highly expressing calcium-binding molecules, *ANXA2* and *SLC3A1*. Immature astrocyte markers and cell cycle-related genes were highly expressed among HD-enriched cluster 3. Several ECM-related genes, including collagens and fibronectin, showed increased expression in HD-enriched cluster 5 along with calcium-binding and adhesion-related genes, which were also enriched in HD-enriched clusters 1 and 2. HD-enriched cluster 8 was the smallest cluster (493 nuclei; 1.4% of total nuclei sequenced) and had a gene expression profile consistent with astroglial progenitors. Non-genotype enriched cluster 7 had a relatively high expression of DNA repair and transcription-related genes, like *FANCA* and *EZH2*. Overall, control states were enriched for astrocyte functions involved in neural support and neuroinflammation, while HD states were enriched for adhesion, ECM, and immature markers. These data highlight both a loss of normal astrocyte functional support, potential developmental impairments, and novel ECM and adhesion-related alterations in HD iAstros. Finally, there is complete loss in HD iAstros of a control state having high expression of *NEAT1,* a lncRNA regulator of transcription that has previously been implicated in HD (Cheng et al., 2018; Sunwoo et al., 2017) and is involved in altered neural cellular development and neural cell damage (Hirose et al., 2014; Katsel et al., 2019).

To further explore the functional relevance of the iAstro clusters to biological pathways, Ingenuity Pathway Analysis (IPA) pathway enrichment was performed on the genes differentially expressed between clusters. The top 2 most significant pathways per cluster were compared to assess the transcriptional signatures unique to each iAstro cluster (**Figure 2E**). Hepatic Fibrosis/Hepatic Stellate Cell Activation was a pathway among the top two significantly enriched pathways in all iAstro clusters except 3 and 6. Control-enriched cluster 0 had significant inhibition of Kinetochore Metaphase Signaling (z=-3.16, p<3e-9), composed of cyclin, centromere, and histone-related molecules (*CCNB1, CDK1, CENPE, CENPK, H2AX, H2AZ1),* which was activated in the immature HD-enriched cluster 3 (z=4.64, p<2e-43). Control-enriched cluster 4 was most significantly enriched in Hepatic Fibrosis/Hepatic Stellate Cell Activation (z=0, p<1e-5) largely due to the downregulation of several collagens (*COL1A1, COL4A1, COL4A2)*. Control-enriched cluster 6 had a significant activation of Neuroinflammatory Signaling molecules (z=3.74, p<6e-14), like HLA transcripts and *SLC1A2*. HD-enriched iAstro cluster 1 had significant enrichment of Antigen Presentation Pathway (z=0, p<3e-7) due to the expression of RNA transcripts that encode various HLA proteins (*HLA-A, HLA-C, HLA-DPA1, HLA-DRB5, HLA-E*). The significant upregulation of several collagen transcripts (*COL11A1, COL18A1, COL1A2, COL27A1, COL3A1, COL4A1, COL4A2, COL6A2*) caused HD-enriched cluster 2 to be enriched in GP6 Signaling molecules (z=0, p<7e-8). HD-enriched iAstro cluster 3 had a significant activation of mitosis-related pathways, like Kinetochore Metaphase Signaling Pathway (z=4.65, p<2e-43) and Cell Cycle Control of Chromosomal Regulation (z=4, p<4e-19) with the upregulation of cyclin-related genes, like *CDK1*, that suggests a mitotic activation of HD iAstros. The dysregulation of integrins (*ITGA11, ITGA6, ITGB1, ITGB4, ITGB8*) and tubulins (*TUBA1A, TUBA1C, TUBB2A, TUBB2B, TUBB6*) induced enrichment of Axonal Guidance Signaling in cluster 5 (z=0, p<3e-16) and 8 (z=0, p<1e-6), which was comprised of 81% HD iAstros. Caveolar-mediated Endocytosis Signaling was significantly enriched (z=0, p<4e7) in cluster 7, which highly expressed integrin beta subunits. These nine iAstro clusters comprise unique transcriptional human astrocytic signatures that provide a glimpse into functionally relevant states of homeostatic and HD astrocytes.

To directly investigate genotype differences between HD and control iAstros, the top 5 significantly activated and top 5 significantly inhibited pathways by genotype were plotted by activation score (**Figure 2F**). Eukaryotic Initiation Factor 2 (EIF2) signaling (z=5.47, p<6e-41) and Actin Signaling Pathways (z=3.68, p<6e-14) were the most significantly activated pathways in HD iAstros, which contained overlapping actin-related genes (*ACTA2, ACTB, ACTC1, ACTG2*). Glutamate Receptor (z=-2.24, p<0.001), Endocannabinoid Neuronal Synapse (z=-2.71, p<0.04), and Neuropathic Pain Signaling Pathways (z=-3, p<0.05) were among the inhibited pathways in HD iAstros and contained genes encoding multiple glutamate receptors, *GRIA1-2* and *GRIN2A-B*. Additional inflammatory-related signaling pathways were among the most significantly inhibited, including Systemic Lupus Erythematosus in T Cell Signaling (z=-2.65, p<0.002) and Coronavirus Pathogenesis Pathways (z=-3.78, p<1e-15) with dysregulation of transcripts in common encoding HLA proteins and ribosome subunits.

Together, HD iAstro gene expression changes by cluster and by genotype highlight an inhibited glutamate receptor signaling state and unique astrocyte cell states, such as actin cytoskeletal signaling activation and immature astrocytes, that may contribute to loss of neural support and altered morphology or motility.

### Defining HD Striatal and Cortical Mouse Astrocyte Cell States

Human HD iAstros show shifts in cell state compositions, including complete loss of a state seen in control iAstros (cluster 4), that represents mature astrocytes with decreased expression of ECM related genes and increased expression of NEAT1. We noticed a shift in the composition of HD iAstros that indicated an abundance of immature and progenitor like cells before and after GLAST+ sorting, these differences may represent deficits in astrogliogenesis which may lead to the functional state changes seen above. We next compared if similar state changes occur in an *in vivo* animal model of HD, and whether species-specific states exist. To assess the transcriptional cell states of striatal and cortical mouse astrocytes within a well-established system that recapitulates transcriptome changes in human HD tissue (Hodges et al., 2006), we carried out snRNA-seq on striatal and cortical brain regions from the rapidly progressing R6/2 HD model, generated through the transgenic overexpression of the first exon of human HTT, and non-transgenic (NT) mice at ages 8-weeks, symptomatic, and 12-weeks, highly symptomatic (Mangiarini et al., 1996) (**Figure 3A** & **4A**).

**Figure 3.**
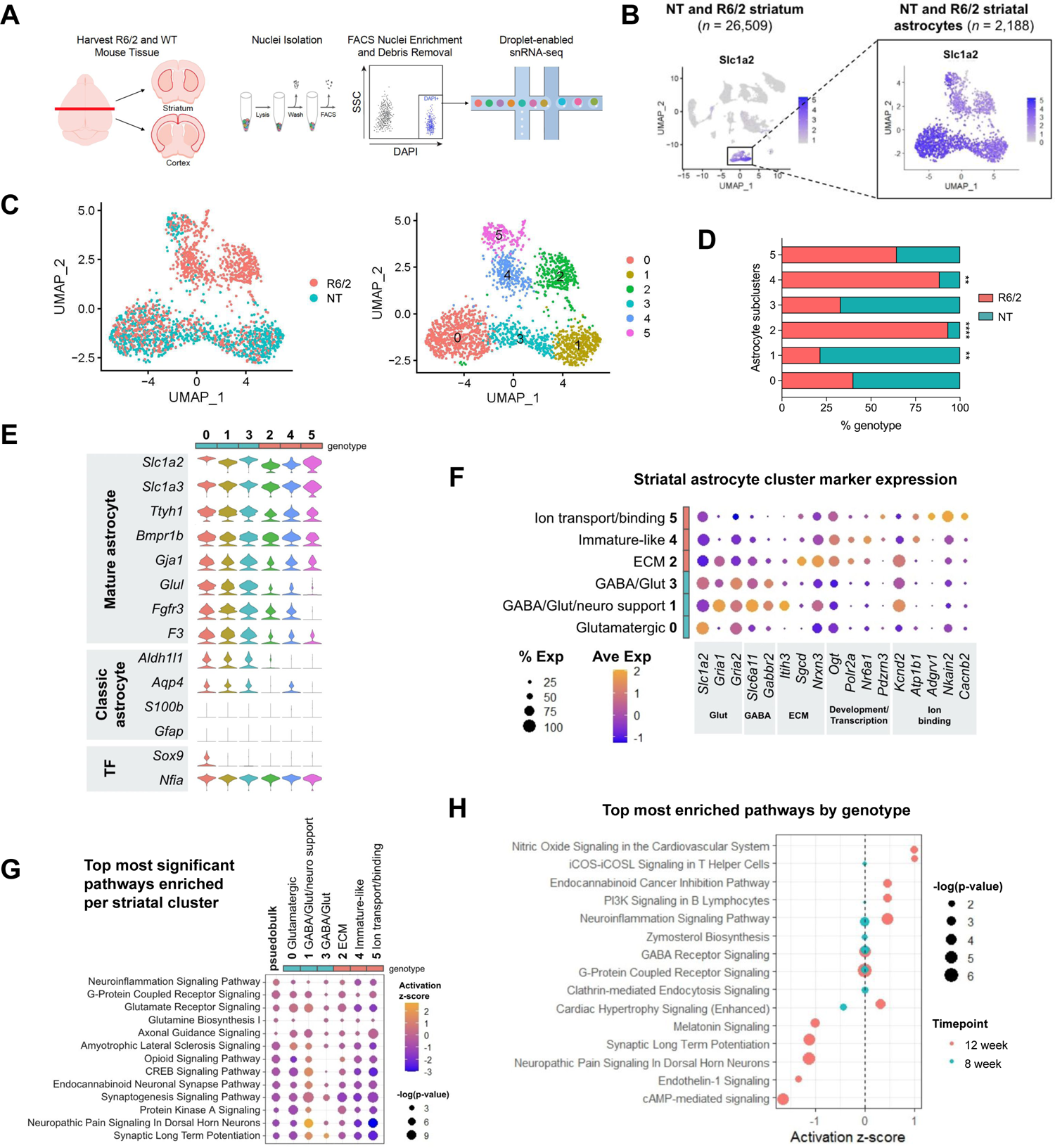
Striatal R6/2 astrocytes exhibit immature and decreased synaptogenesis cell states. (**A**) Experimental workflow for snRNA-seq analysis where striatal and cortical tissue was harvested from NT and R6/2 mice, dissociated, then underwent single-nuclei isolation and FACS separation prior to droplet-enabled snRNA-seq. (**B**) UMAP of all 12-week striatal snRNA-seq libraries performed on aggregated peak matrix. The astrocyte cluster was identified by expression of *Slc1a2*. (**C**) UMAP subset analysis of astrocyte cluster identified several astrocyte subclusters across R6/2 and NT. (**D**) Genotype composition of cells across astrocyte subclusters (two-way ANOVA, Bonferroni multiple comparisons test performed on number of nuclei/mouse within each subcluster adjusted p-values: cluster 0 p=0.0809, cluster 1 **p=0.0012, cluster 2 ****p<0.0001, cluster 3 p=0.5160, cluster 4 **p=0.0057, cluster 5 ns; n=3 mice per genotype per timepoint). (**E**) Astrocyte marker genes across each subcluster shows a decrease in maturation gene expression in several striatal R6/2-enriched subclusters. Genotype enrichment was defined by >60% of cluster, gray genotype represents 40-60%. (**F**) Top cluster markers for each cluster for cell state classification. (**G**) Top significant pathways across all striatal astrocyte clusters used to classify cell state signatures. Psuedobulk genotype pathway enrichment using DEGs from all 12-week R6/2 striatal astrocytes compared to all 12-week NT striatal astrocytes is included for comparison. (**H**) Top most significantly enriched pathways by age for 8- and 12-week striatal R6/2 astrocytes.

At the 12-week time point, we identified 13 and 16 distinct clusters from 26,509 striatal and 25,237 cortical nuclei sequenced, respectively. The *Slc1a2* astrocyte marker gene that encodes GLT1, a glutamate transporter that clears the excitatory neurotransmitter from the extracellular space at synapses, was used as an astrocyte marker and expression was plotted across all cells to identify the astrocyte cluster for each dataset (**Figure 3B** & **4B**). The cluster with the highest *Slc1a2* expression was subset to investigate R6/2 astrocyte-specific dysregulation via sub-clustering and further transcriptomic analyses. Due to a small population of cells with relatively low levels of astrocyte markers and high expression of vascular markers *Pdgfrb* and *Flt1* (Zhang et al., 2014), additional sub-setting was necessary before sub-clustering the cortical astrocyte cluster (**Figure S4**). Upon sub-clustering all 12-week R6/2 astrocyte clusters, we identified six striatal astrocyte clusters and five cortical astrocyte clusters visualized using UMAP (**Figures 3C** & **4B**). Interestingly, the total number of clusters identified were less than the human iAstros which may represent species differences, considering the known complexity of human glia versus mouse glia (Oberheim et al., 2009). Diverse cellular states were represented across each mouse dataset and to identify genotype composition by cluster, the total number of cells per genotype were quantified by cluster. Striatal clusters 2 and 4 were significantly enriched in R6/2 compared to NT, while cortical cluster 0 was significantly enriched in R6/2, indicating these clusters are relatively novel to the R6/2 condition (**Figures 3D** & **4C**). R6/2 striatal astrocytes had a larger number of differentially expressed genes by genotype compared to cortical astrocytes (**Figures S5** & **S6**).

Expression of mature astrocyte marker genes and transcription factors regulating astrogliogenesis were visualized across astrocyte clusters (**Figure 3E** & **4D**). R6/2-enriched striatal astrocyte clusters 2 and 4 as well as R6/2-enriched cortical cluster 0 had a lower expression of several classic astrocyte identity genes, such as *Slc1a2* and Glutamate-Ammonia Ligase (*Glul*), compared to NT-enriched clusters. Similarly, cortical astrocyte clusters 1 and 3 had a lower expression of these astrocyte identity genes compared to cluster 0. Decreased expression of astrocyte markers in R6/2-enriched clusters indicates that dysregulated cell states exist in R6/2 astrocytes that may reflect differences in astrocyte development or a loss of cell identity genes. Regardless, there was a shift in the R6/2 astrocytes towards more immature cell states.

### Inhibited Synaptogenesis-Related Signaling States in R6/2 Striatal Astrocytes

We next molecularly assessed the cell states identified in striatal and cortical R6/2 and NT mouse astrocytes. First, the top highly expressed gene markers per cluster were used to classify clusters into functionally relevant categories (**Figure 3F** & **4E**). Several of the highest expressing genes in NT-enriched striatal clusters 0, 1, 3 and NT-enriched cortical cluster 2 and 1 included astrocyte-enriched glutamate transporter (*Slc1a2) and* glutamate receptors *(Gria1*, *Gria2, Grin2c),* like control-enriched iAstro clusters 0 and 6. NT-enriched striatal clusters 1 and 3 also had high expression of GABA transporter *Slc6a11* and GABA receptor *Gabbr2.* Cortical clusters 1 and 2 had a high expression of an actin-binding gene *Clmn* and a phagocytic receptor *Mertk.* NT-enriched cortical cluster 4 had a high expression of neural regulatory genes (*Anks1b and Nrg3*), axonal/synaptic related genes (*Robo2, Dlg2, and Lrtm4*), and *Kcnip4*, which encodes for a potassium channel interacting protein.

Among the striatal clusters with significantly more R6/2 cells compared to NT, striatal clusters 2 and 4 had the highest expression of developmental-related genes, RNA polymerase (*Polr2a*) and a nuclear receptor involved in neurogenesis (*Nr6a1*). R6/2-enriched striatal cluster 2 also had the highest expression of two ECM-related genes, glycoprotein-related *Sgcd* and cell adhesion molecule *Nrxn3*. R6/2-enriched cortical cluster 0 highly expressed *Ogt*, a glycosyltransferase regulator. Similar to HD-enriched iAstro clusters 1 and 2, R6/2-enriched striatal cluster 5 had the highest expression of sodium, potassium, or calcium ion binding genes, like *Adgrv1, Nkain2,* and *Cacnb2*, except for the potassium channel, *Kcnd2*, which was most highly expressed in striatal clusters 1 and 2. Cortical cluster 3 exhibited high expression of calcium-binding genes, *Adgrv1* and *Cacnb2*, and was not enriched by genotype.

To further explore the potential functional relevance of these striatal astrocyte clusters, we ran pathway enrichment analysis (IPA) on the DEGs for each cluster compared to all other striatal (**Figure 3G**) and cortical astrocytes (**Figure 4F**). NT-enriched striatal cluster 0 had the highest activation of an Amyotrophic Lateral Sclerosis (ALS) Signaling pathway (z=1, p<4e-6) and Glutamate Receptor Signaling (z=0.38, p<1e-8) due to the high expression of overlapping genes that encode proteins responsible for glutamate-glutamine cycling in astrocytes (*Slc1a2, Glul, Gria1, Grid1,* and *Grin2c)*. Also significantly enriched in NT astrocytes, striatal cluster 1 had the highest activation in Neuropathic Pain Signaling (z=2.67, p<3e-8) and Synaptic Long Term Potentiation (z=1.6, p<7e-7), due to the overlap in upregulated genes *Gria1, Gria2,* and *Grin2c*. NT-enriched striatal cluster 3 exhibited significant activation of Synaptic Long Term Potentiation (z=2, p<0.001) and Synaptogenesis Signaling (z=-1.35, p<9e-10), with upregulation of *Gria2* in both pathways. Cortical cluster 4 was largely composed of NT astrocytes and had significant activation of Synaptogenesis Signaling (z=6.97, p<7e-29) and Calcium Signaling (z=6.08, p<3e-13) through upregulation of genes that encode calcium/calmodulin-dependent protein kinases (*Camk2a, Camk3b, Camk4*) and calcium voltage-gated channel subunits (*Cacna1b, Cacna2d1, Cacnb4*).

**Figure 4.**
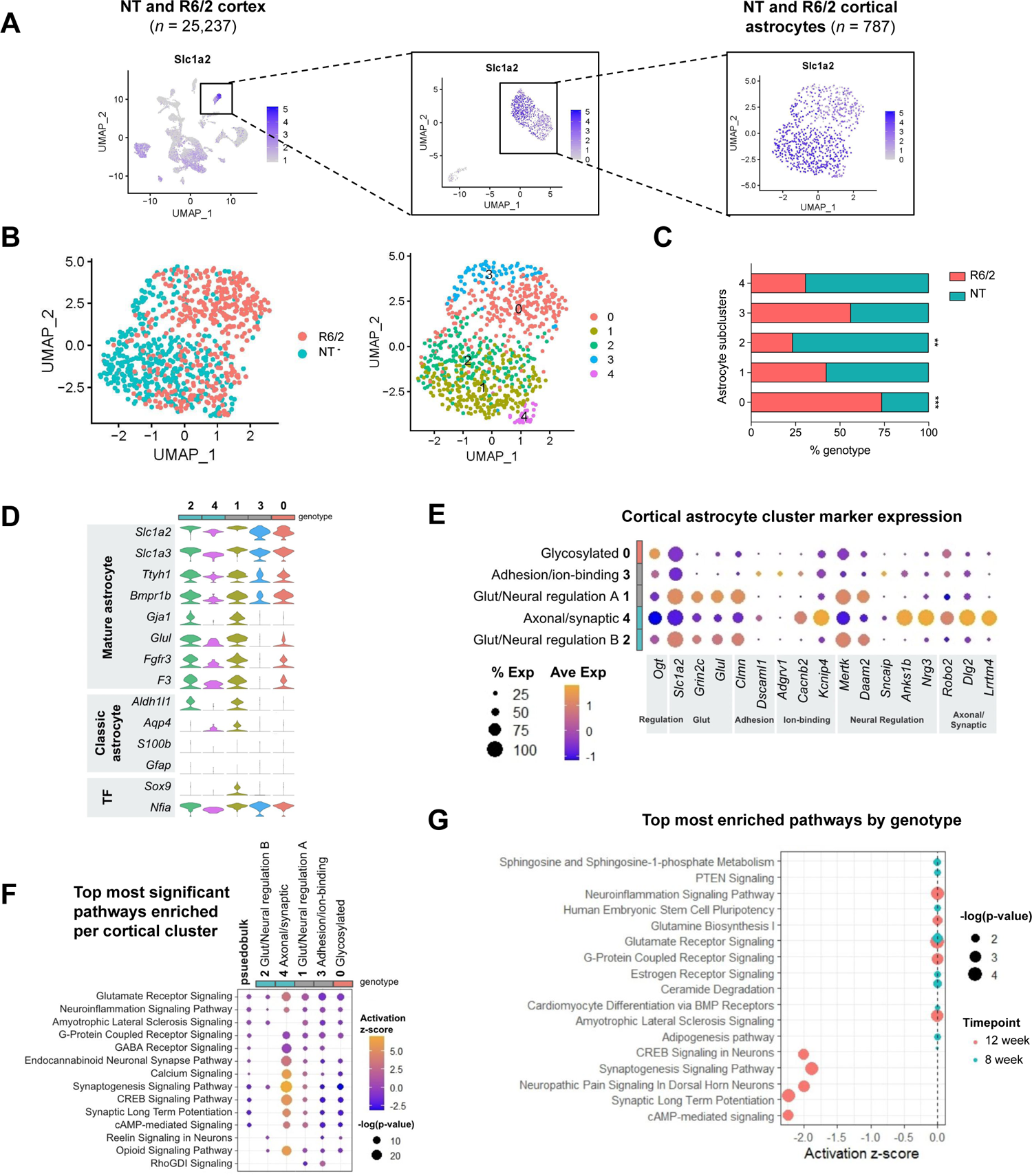
Cortical R6/2 astrocytes exhibit inhibited neuronal homeostatic signaling and decreased astrocyte marker expression. (**A**) UMAP of all 12-week cortical snRNA-seq libraries performed on aggregated peak matrix. The astrocyte cluster was identified by expression of astrocyte marker gene, *Slc1a2*. Additional subclustering was necessary to subset out cortical vascular cells from astrocytes. (**B**) UMAP subset analysis of astrocyte cluster identified several astrocyte subclusters across R6/2 and NT. (**C**) Genotype composition of cells across astrocyte subclusters (two-way ANOVA, Bonferroni multiple comparisons test performed on number of nuclei/mouse within each subcluster adjusted p-values: cluster 0 ***p=0.0004, cluster 1 p=0.8024, cluster 2 **p=0.0064, cluster 3 ns, cluster 4 ns; n=3 mice per genotype per time point). (**D**) Astrocyte marker genes across each subcluster. Genotype enrichment was defined by >60% of cluster, gray genotype represents 40-60%. (**E**) Top cluster markers for each cluster for cell state classification. (**F**) Top significant pathways across all cortical astrocyte clusters used to classify cell state signatures. Psuedobulk genotype pathway enrichment using DEGs from all 12-week R6/2 cortical astrocytes compared to all 12-week NT cortical astrocytes is included for comparison (**G**) Top most significantly enriched pathways by age for 8- and 12-week cortical R6/2 astrocytes.

NT-enriched cortical clusters 1 and non-genotype-enriched 2 were significantly enriched in Glutamate Receptor Signaling due to expression of *Grin2c* and *Slc1a2*, with predicted activation in cluster 1 (z=1, p<5e-8) but no predicted activation for cluster 2 (z=0, p<0.002). Cortical cluster 1 also exhibited a predicted activation of Synaptogenesis Signaling (z=2.83, p<0.0006). Glutamate Receptor Signaling was the most significantly enriched pathway for cortical cluster 3, with a predicted inhibition (z=-1.34, p<4e-11), like HD iAstros. Additionally, Rho GDP-dissociation inhibitor (RhoGDI) Signaling was predicted to be most activated and Synaptogenesis Signaling was the most inhibited predicted pathway for non-genotype-enriched cortical cluster 3.

Among the striatal astrocyte clusters significantly enriched with R6/2 cells, the most significant pathways across striatal clusters 2 and 4 were Synaptogenesis Signaling (cluster 2: z=-1.35, p<9e-10; cluster 4: z=-1.81, p<2e-7) and Glutamate Receptor Signaling (cluster 2: z=0, p<3e-7; cluster 4: z=-1.34, p<9e-7) specifically due to the downregulation of genes associated with glutamate receptors, like *Gria1* and *Gria2.* Neuropathic pain (z=-3, p<1e-9) was among the most inhibited pathways for striatal R6/2-enriched cluster 5, as well as Synaptic Long Term Potentiation (z=-2.36, p<8e-10) and cAMP response element-binding protein (CREB) Signaling (z=-2.36, p<1e-8). R6/2-enriched cortical cluster 0 was most significantly enriched in Glutamate Receptor Signaling (z=-1, p<7e-9), and Synaptogenesis signaling pathway was predicted to be the most inhibited pathway (z=-3.16, p<2e-6).

Overall, the data suggest a loss of glutamate signaling and synaptogenesis-related pathways across R6/2 astrocytes, and activation of ECM and immature-like astrocyte states, similar to human HD patient iAstros. These data highlight similarities with the human HD iAstros that suggest maturation impairments, with a particular focus on glutamate receptor and synaptogenesis-related dysregulation.

### HD Mouse Astrocyte Temporal and Brain Region Comparisons

We next examined transcriptional changes that occur in R6/2 astrocytes with progressive disease. We assessed the activation scores of the top ten most significant pathways enriched in 8-week mouse astrocyte signaling states and 12-week mouse astrocyte signaling states compared to age-matched NT mouse striatal astrocytes (**Figure 3H**) (Luthi-Carter et al., 2002). As expected, there was overlap of many relevant pathways between the 8- and 12-week striatal R6/2 astrocytes, with the 12-week astrocytes exhibiting increased predicted activation of iCOS-iCOSL Signaling in T Helper Cells (12-week: z=1, p<0.008; 8-week: z=0, p<0.032), PI3K Signaling in B Lymphocytes (12-week: z=0.447, p<0.003; 8-week: z=0, p<0.048), and Neuroinflammation Signaling (12-week: z=0.447, p<5e-5; 8-week: z=0, p<0.001). *Chuk*, which encodes an inhibitor of the transcription factor NFκB complex, was an overlapping upregulated gene across the three inflammatory pathways. The most significantly inhibited pathway in 12-week striatal astrocytes was cAMP-mediated Signaling (z=-1.67, p<3e-5). Interestingly, Cardiac Hypertrophy Signaling was the most significantly inhibited pathway predicted in the 8-week striatal R6/2 astrocytes (z=-0.447, p<0.008) due to expression of genes such as *Camk2g*, *Chuk*, and *Fgf14*, but was predicted to be activated in 12-week striatal R6/2 astrocytes (z=0.302, p<6e-4). In general, the most significantly enriched pathways in 8-week striatal astrocytes had no predicted activation or inhibition. By 12 weeks, R6/2 astrocytes had more severe predicted activation or inhibition of these overlapping pathways that correlates with increased severity in motor phenotypes (Luthi-Carter et al., 2002), suggesting increased astrocyte dysfunction with disease progression. The overlap in top pathways between 8- and 12-week with progressive dysregulation of similar genes between 8 and 12 weeks indicates that at these two timepoints, R6/2 astrocytes are similarly dysregulated in symptomatic animals. Taken together, R6/2 astrocytes exhibited inhibition of neuronal homeostatic signaling pathways, suggesting how mHTT expressing striatal astrocytes could contribute to neuronal dysregulation in R6/2 mice.

Temporal changes occurring in R6/2 cortical astrocytes were also compared for cortical astrocytes (**Figure 4G**). The most significantly enriched pathways in 12-week cortical astrocytes were predicted to be inhibited or had no activation scores. Among those were Glutamate Receptor Signaling (z=0, p<8e-5), Synaptic Long Term Potentiation (z=-2.236, p<2e-4), and Neuroinflammation Signaling Pathway (z=0, p<2e-4), with glutamate receptor *Grin2c* being a common downregulated gene across the three pathways. The most significantly enriched pathway in 8-week cortical astrocytes was Glutamate Receptor Signaling (z=0, p<0.004), which was also significantly enriched in 12-week cortical astrocytes due to downregulation of related genes (*Glul, Gria2, Grin2c,* and *Slc1a2*). The 8-week cortical astrocyte pathways had no predicted activation or inhibition (z=0), like 8-week striatal astrocytes. These findings further demonstrate dysregulated signaling in R6/2 astrocytes related to neuronal homeostasis, particularly Glutamate Receptor Signaling that overlap with HD iAstros and increase in severity with disease progression, although diversity in cell states was less severe in R6/2 cortical astrocytes compared to R6/2 striatal astrocytes.

To determine the impact of mHTT on R6/2 astrocyte cell states by brain region, we directly compared striatal and cortical astrocyte pathway enrichment performed on genotype DEGs from both timepoints (**Figure 5A** and **S4**). R6/2 12-week striatal and cortical astrocytes showed significant inhibition of neural regulatory-related signaling, including Synaptic Long Term Potentiation, Synaptogenesis, and CREB Signaling in Neurons. The most significant pathways enriched in R6/2 8-week striatal and cortical astrocytes had low or no predicted activation scores, but the individual pathways were more unique to each brain region. When evaluating each region, striatal 8-week top pathways included Neuroinflammation and GABA receptor signaling, while cortical 8-week top pathways were Glutamate Receptor Signaling, Ceramide Degradation, and Sphingosine-1-phosphate (S1P) metabolism; this highlights the unique cellular processes in metabolic regulation across these two brain regions during this stage in HD pathogenesis that may drive changes at later stages. There were many overlapping significant pathways relating to Synaptogenesis Signaling in 12-week striatal and cortical timepoints but had increased predicted inhibition in cortical astrocytes. These shared and unique pathways across astrocytes in age and brain region highlight the unique astrocyte cell states that may contribute to R6/2 pathogenesis across these two highly affected brain regions.

**Figure 5.**
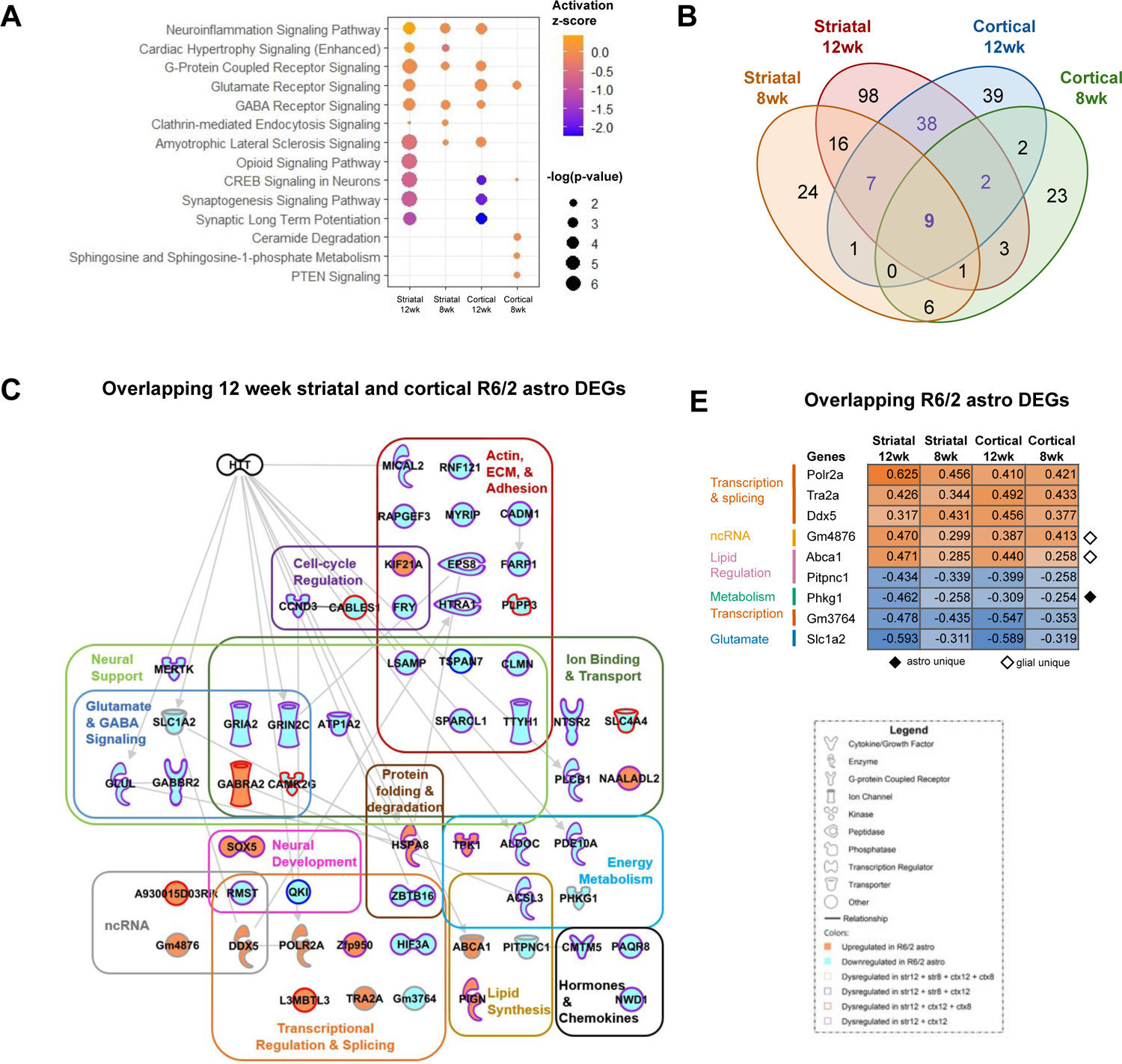
Shared R6/2 astrocyte dysregulation across brain regions and ages highlights neuronal homeostasis dysregulation. (A) Top pathways dysregulated across all four R/2 datasets (**B**) Venn diagrams of significant differentially expressed genes (DEGs) in R6/2 mouse astrocytes across time points and brain region (Exact hypergeometric probability calculated using 56 DEG overlap for 12-week striatum [174] vs 12-week cortex [98] ****p<8e-113, 33 DEG overlap for 12-week striatum [174] vs 8-week striatum [64] ***p<7e-64, 13 DEG overlap for 12-week cortex [98] vs 8-week cortex [46] *p<8e-25, 16 DEG overlap for 8-week striatum [64] vs 8-week cortex [46] **p<3e-35 using 46,206 as the number of genes with nucleotide sequence data in the mouse genome database). (**C**) Top most significant canonical pathways across brain regions and ages. Overlap of 12-week striatal and cortical astrocyte DEGs highlight pathway dysregulation in R6/2 astrocytes across both brain regions in symptomatic mice. (**E**) The nine overlapping DEGs across all datasets. Orange represents upregulated DEG and blue represents downregulated DEG. Solid black diamond denotes a DEG unique to only astrocytes, while a white diamond denotes a DEG unique to astrocytes, oligodendrocytes, and OPCs.

As the overlapping pathways across time points and brain regions suggested, there was significant astrocyte genotype DEG overlap (**Figure 5B**). The set of 56 overlapping 12-week striatal and cortical DEGs were genes all dysregulated in the same direction and may represent common signatures in astrocytes independent of region (**Figure 5C**). Individual examination of each of these genes was performed to uncover commonalities contributing to transcriptional dysregulation in R6/2 mouse astrocytes at this severe disease stage. *Gria2* and *Grin2c* were present in a majority of the overlapping dysregulated signaling pathways related to neuronal support, glutamate receptor signaling, and ion binding, highlighting how aberrant R6/2 astrocytes affect the regulation of neuronal homeostasis. There were also several genes related to both neural support and extracellular matrix/adhesion-related signaling, such as *Sparcl1, Ttyh1*, and *Tspan7*. Together, this dysregulated astrocyte signaling at 12 weeks may highlight two major dysfunctions occurring across R6/2 astrocytes and how it affects the cells they are meant to support, therefore contributing to pathogenesis.

To further assess the common dysregulation across R6/2 astrocytes, we examined the nine genotype DEGs that were common across astrocytes from both brain regions and time points. These genes all had very similar expression levels that often became more severe with age, with striatal 12-week astrocytes typically having the most extreme expression of these nine common DEGs (**Figure 5E**). Exceptions included several splicing and transcription-related genes (*Gm3764, Tra2a, Ddx5*) that had the highest fold change in DEGs from cortical 12-week R6/2 astrocytes compared to cortical NT astrocytes. A non-coding RNA (*Gm4876*) and a lipid regulator (*Abca1*) were DEGs unique to only glial cell types (astrocytes, oligodendrocytes, and/or oligodendrocyte progenitor cells) in the larger R6/2 snRNA-seq dataset. Among these overlapping astrocyte DEGs, the downregulation of the glycogen phosphorylase activator, *Phkg1,* was unique to only astrocytes compared to other R6/2 cell types and highlights the potential dysregulation of glycogen breakdown in R6/2 astrocytes. These common DEGs may represent core processes, metabolic and transcriptional, that become dysregulated at an early stage of HD pathogenesis and give rise to other changes seen at 8 and 12 weeks in the R6/2 mice. The changes in these metabolic and transcriptional processes may lead to the altered signaling described above related to synaptogenesis, glutamate signaling, calcium signaling, and inflammation.

### Inhibition of Glutamate Signaling in HD iAstros and R6/2 astrocytes

A significantly inhibited glutamate receptor signaling cell state was identified in HD-enriched R6/2 and iAstro clusters. The presence of these states in both the mouse and iPSC model suggest that this is a core feature of HD astrocytes and likely arises intrinsically from the expression of mHTT. Genes that encode glutamate transporters (*SLC1A2*, *SLC1A3*) and glutamate receptors (*GRIA1, GRIA2, GRIN2A*) were significantly downregulated across both models of HD astrocytes (**Figure 6A & B**). To confirm the intrinsic loss of glutamate signaling in HD astrocytes, we investigated protein levels of a representative glutamate receptor 1 (GluR1), encoded by *GRIA1;* Western analysis for GluR1 was performed on HD and control iAstros and showed significantly decreased GluR1 in HD iAstros compared to control iAstros (**Figure 6C**). Due to the common down regulation seen in both mouse and human HD astrocytes in this study and in previously published results, we next assessed if our HD iAstros showed a functional impairment. HD iAstros demonstrated decreased functional uptake of glutamate when exogenous glutamate was added *in vitro* (**Figure 6D**). Since glutamate signaling, and specifically downregulation of glutamate transporters and receptors, was observed in both models with the iAstros showing functional downregulation, we suspect that this is a cell-autonomously downregulated signaling pathway in HD astrocytes. The sorted iAstros are not exposed to signaling from neurons, and recapitulation of this alteration provides evidence for cell autonomous effects. Inhibition of the uptake of glutamate, can influence HD pathogenesis by contributing to glutamate neurotoxicity.

**Figure 6.**
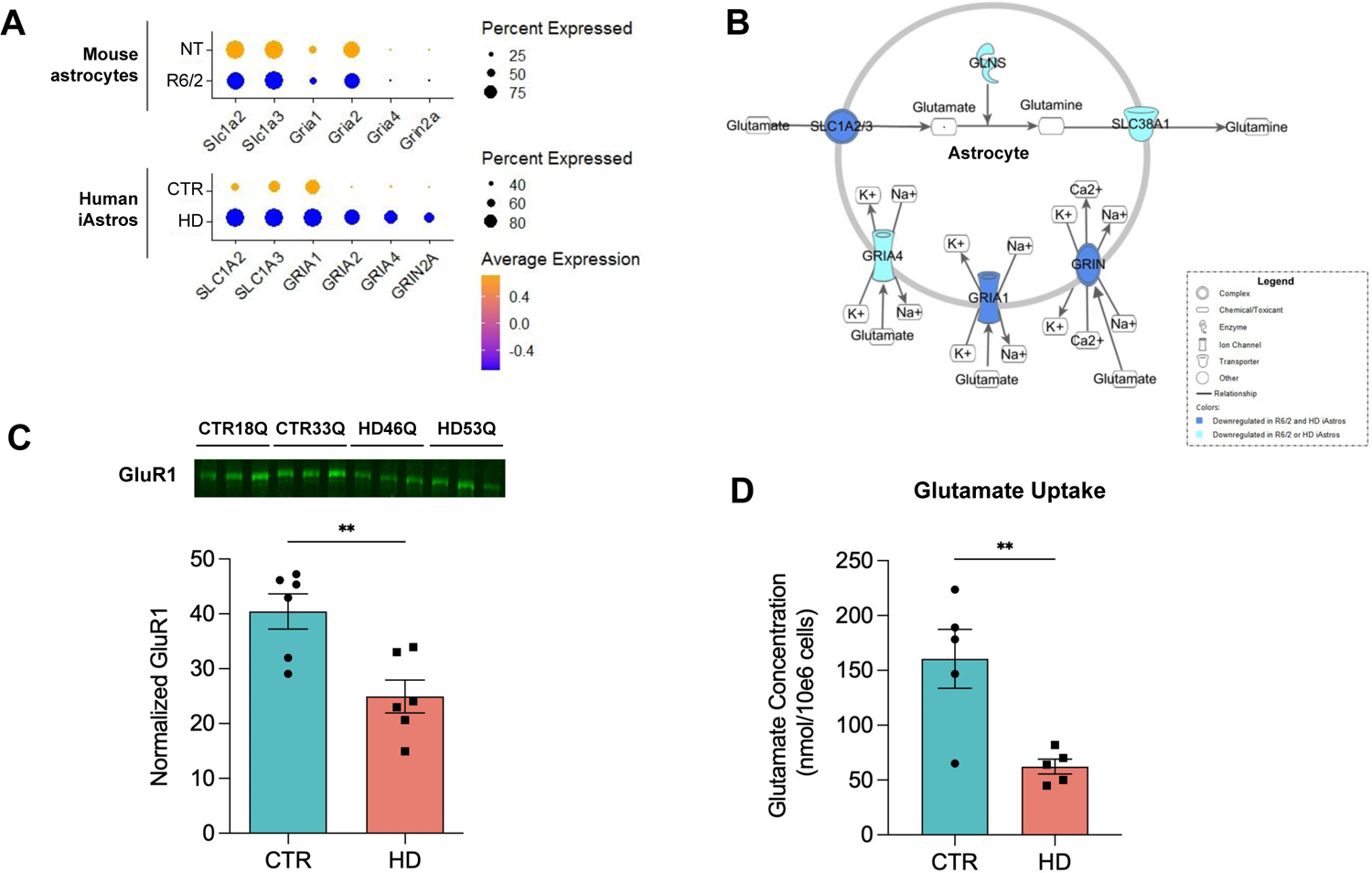
HD astrocyte inhibited glutamate receptor signaling in R6/2 and HD iAstros. (**A**) Significantly downregulated molecules in glutamate receptor signaling pathway across R6/2 and HD iAstros. (**B**) Glutamate receptor signaling network R6/2 and HD iAstro snRNA-seq shows significant downregulation of glutamate-related molecules across multiple models of HD astrocytes. (**C**) GluR1 Western blot and quantification demonstrates significant decreased GluR1 in HD iAstros (unpaired t-test, **p=0.0054, n=3 biological [differentiation] replicates/line, 2 lines/genotype). (**D**) HD iAstros uptake significantly less glutamate than control (CTR) iAstros (unpaired t-test, **p=0.0075, n=2-3 biological [differentiation] replicates/line, 2 lines/genotype). All error bars indicate ± standard error mean.

### Activation of ECM and Actin in HD iAstros

ECM and actin cytoskeletal signaling were processes uniquely activated in a subcluster of HD iAstros which were not identified in R6/2 astrocytes. While there were several actin, ECM, and adhesion-related DEGs (e.g. *CADM1, CLMN, MICAL1*) present in overlapping 12-week striatal and cortical R6/2 astrocytes (**Figure 5C**), this did not trigger significant enrichment of specific actin or ECM signaling pathways, unlike HD iAstros. We investigated the genes contributing to ECM and actin cytoskeletal signaling activation in the HD and control iAstro cell lines to determine the genes controlling these dysregulated pathways (**Figure 7A**). Genes related to extracellular signaling (*ITGB1* and *ITGA3*) and those regulating actin dynamics (*ACTN1, Talin, ERM*), actin polymerization (*F-actin, PFN, and ARP2/3*), and overall cytoskeletal reorganization (*MYL and Myosin*) were all significantly upregulated in HD iAstros (**Figure 7B**). Given the potential disruption of the actin cytoskeleton, we investigated whether filamentous actin (F-actin) protein expression was increased in our HD iAstros as a surrogate readout. Phalloidin in-cell westerns were performed on HD and control iAstros and showed that HD iAstros had significantly increased expression of F-actin compared to control iAstros (**Figure 7C**). Additionally, HD iAstros exhibited increased levels of the F-actin crosslinking protein, Actinin (ACTN), compared to control iAstros (**Figure 7D**), suggesting increased actin cytoskeletal scaffolding as a gain of cell state in HD iAstros. Taken together, ECM and actin cytoskeletal signaling are cell-autonomous and potentially early activated signaling pathways unique to patient-derived HD astrocytes that may contribute to morphological changes or increased adhesion in HD astrocytes.

**Figure 7.**
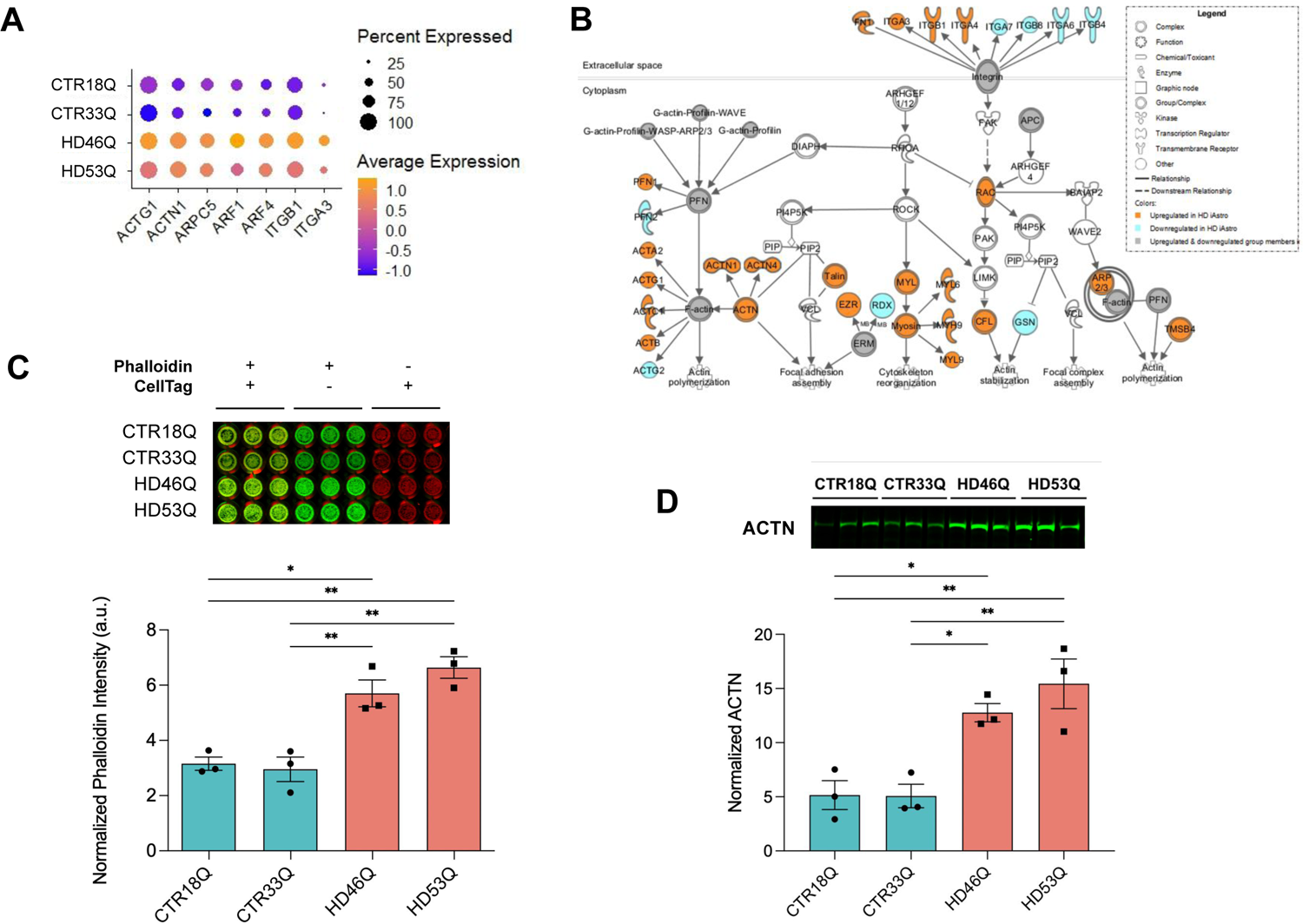
Actin Cytoskeletal and Integrin Signaling Activation in Human HD iAstros. (**A**) Significantly downregulated molecules in integrin and actin cytoskeletal signaling pathways HD and control iAstro lines. (**B**) Pathway or network from snRNAseq including ITGs and Actin. (**C**) Phalloidin in-cell Western blot and quantification shows significant upregulation of F-actin in HD iAstro cell lines (one-way ANOVA, n=3 biological [differentiation] replicates/line; 18Qv46Q: *p<0.05; 33Qv46Q, 18Qv53Q, 33Qv53Q: **p<0.01; 18Qv33Q, 46Qv53Q: ns). (**D**). ACTN western blot and quantification shows significant increased ACTN in HD iAstros (one-way ANOVA, n=3 biological [differentiation] replicates/line; 18Qv46Q, 33Qv46Q: *p<0.05; 18Qv53Q, 33Qv53Q: **p<0.01; 18Qv33Q, 46Qv53Q: ns). All error bars indicate ± standard error mean.

### Aberrant Astrogliogenesis Transcriptional Regulators in HD Astrocytes

To investigate the potential regulatory mechanisms controlling human HD astrocyte dysregulation, we used genotype DEGs in HD iAstros to predict transcription factor regulation enrichment analysis by Chromatin immunoprecipitation-X Enrichment Analysis (ChEA) (Chen et al., 2013; Kuleshov et al., 2016) (**Figure 8A**). The top predicted transcriptional regulator was an important regulator of astrogliogenesis, ATF3, followed by transcriptional repressor ZNF217 (**Figure 8A**). To validate this finding, we looked at protein levels of ATF3, which were significantly decreased in HD compared to control iAstros, suggesting that aberrant HD astrogliogenesis may be driven by known transcriptional regulators of this process (**Figure 8B**). Furthermore, SOX9, a transcriptional regulator of glial development and early astrogliogenesis, which was significantly downregulated in HD iAstros, also had significantly decreased protein levels (**Figure 8C**). The neural stem cell factor SOX2, was also enriched. YAP1 of the HIPPO signaling pathway that has an essential role in regeneration and self-renewal, was another significantly enriched factor in HD iAstros that has previously been implicated in HD and shown to interact with Htt (Mueller et al., 2018). Overall, HD astrocyte transcriptionally dysregulated cell states and predicted alterations in transcription factors may implicate an astrogliogenesis deficit that induces astrocyte dysfunctions in glutamate signaling and activation of ECM and actin cytoskeletal signaling to potentially contribute to HD pathogenesis via altered glutamate uptake and cellular motility, respectively.

**Figure 8.**
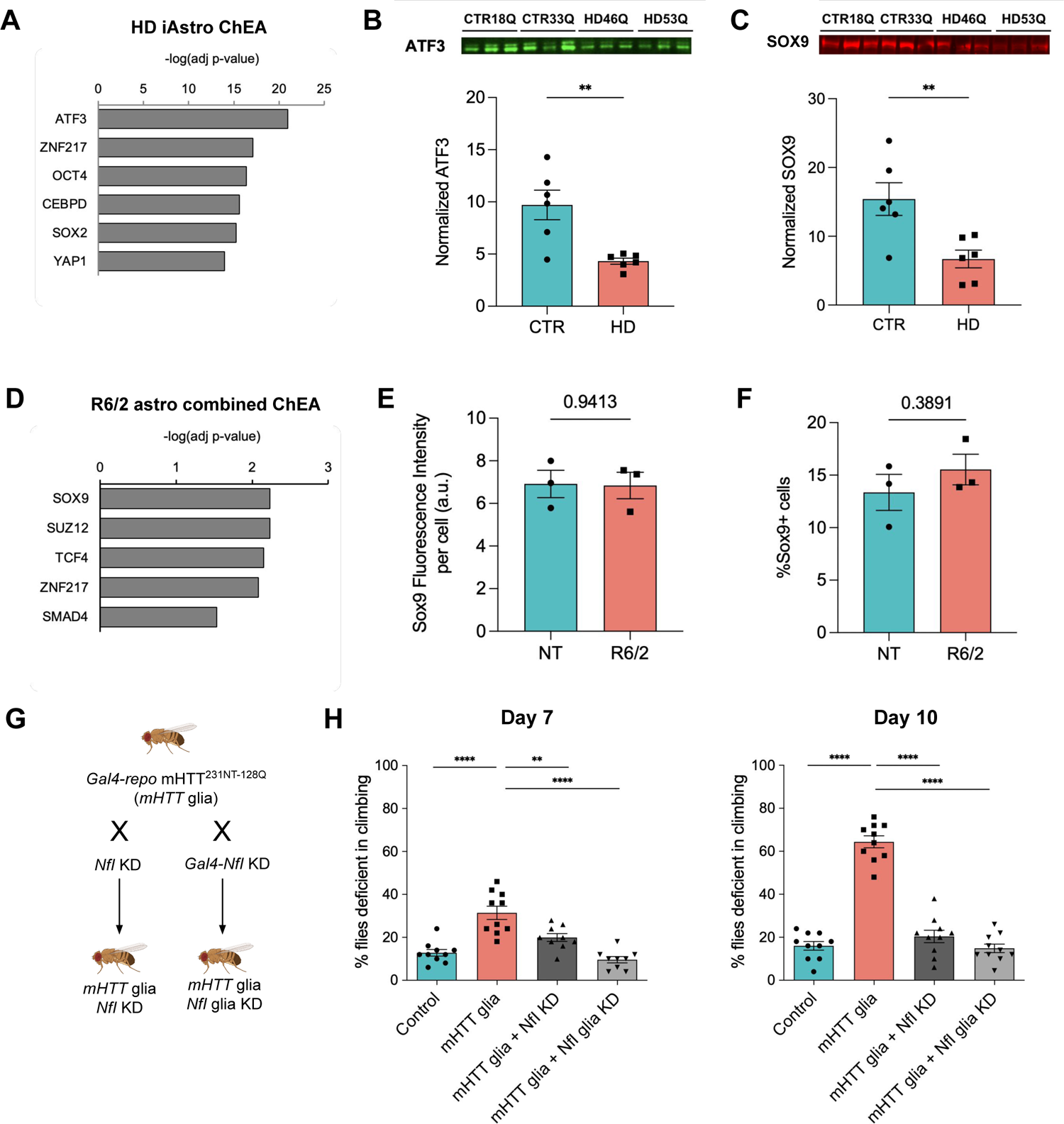
Transcriptional regulation of HD astrocytes. (**A**) Predicted transcription factor regulation in HD iAstros. (**B-C**) ATF3 (B) and SOX9 (C) western blot and quantification shows significant decreased ATF3 (unpaired t-test, **p=0.004, n=3 biological [differentiation] replicates/line, 2 lines/genotype) and SOX9 (unpaired t-test, **p=0.009, n=3 biological [differentiation] replicates/line, 2 lines/genotype) in HD iAstros. (**D**) Overlap of 12-week striatal and cortical astrocyte DEGs predicted transcription factor regulation in R6/2 astrocytes across both brain regions in symptomatic mice. (**E, F**) Immunohistochemistry in nontransgenic (NT) or R6/2 mouse striatum show no significant difference in intensity or number of astrocytes (GFAP-positive or S100β-positive cells) that express SOX9. (**G**) An HD fragment Drosophila model (HTT^231NT128Q^) was crossed with lines that reduce expression of NfI, the fly ortholog to human NFIA, in glia using the Gal4 system. Fly orthologs are listed on the X-axis with the human gene names listed paranthetically. Numbers represent BDSC stock numbers. (**H**) A climbing assay was performed at day 7 and day 10. When mHTT was expressed in glia (grey dots), flies demonstrate a temporal increase in climbing deficiency. When fly lines that reduce expression of target genes were crossed with flies that express mHTT only in glia (black dots), the climbing deficit was largely suppressed. Unpaired t-tests were performed comparing each line to the mHTT-expressing line within each dataset (****p<0.0001, **p=0.01; n=10 biological replicates, each containing 9-10 animals with n=5 consecutive trials/replica/day). All error bars indicate ± standard error mean.

To identify possible transcriptional regulators in R6/2 astrocytes, we conducted transcription factor enrichment analysis (Kuleshov et al., 2016) using the 56 common 12-week DEGs (**Figure 5C**). The ChEA database showed glial developmental transcription factor, SOX9, as the most significant transcription factor enriched, suggesting a role for this complex in altered gene expression in R6/2 mouse astrocytes (**Figure 8D**). TCF4, a transcription factor involved in nervous system development, and SUZ12, a member of the polycomb repressive complex 2 (PRC2), were also enriched, suggesting roles for these complexes in repressing astrogliosis genes in HD (**Figure 8D**), that have also been implicated in HD neurons (Seong et al., 2010), endothelial genes in HD BMECs (Ferrari Bardile et al., 2019; Lim et al., 2017), and HD oligodendroglia (Ferrari Bardile et al., 2019). Another predicted transcriptional regulator in R6/2 astrocytes was the transcriptional repressor ZNF217, also present in HD iAstro analysis (**Figure 8A** & **8D**). SOX9 protein expression was evaluated in the striatum of 12-week-old R6/2 mice; however, there was no significant change in expression levels or number of SOX9-expressing astrocytes (GFAP-positive or S100b-positive cells) (**Figure 8E** & **8F**). For further investigation into transcription factor perturbation in R6/2 mouse astrocytes, Enrichr Transcription Factor Database was queried (**Figure S5B**). Hes Family BHLH Transcription Factor 1 (HES1) overexpression was implicated as the most significant perturbation, further implicating transcriptional repression of nervous system development in R6/2. Alterations in ASCL1 and MECP2, regulators of neural development, were also predicted to contribute to common dysregulated gene sets in R6/2 striatal and cortical astrocytes. NFIA, a transcription factor that controls astrogliogenesis, was identified as being deficient, indicating a potential mechanism for altered astrocyte development in R6/2. These findings highlight potential mechanisms in astrogliogenesis and neural developmental dysregulation contributing to downstream pathway dysregulation in mHTT-expressing astrocytes—a similar finding to our iAstro model of HD.

### Modulation of Astrogliogenesis Regulator NFIA/Nfl in HD Drosophila Glia

We next evaluated if perturbation of an astrogliogenesis transcriptional regulator can rescue HD astrocyte-induced deficits *in vivo*. Given the ease of genetic manipulation, short life cycle, and simple functional assays used to assess complex glial biology and previous data showing a deficit in glia in HD model flies (Al-Ramahi et al., 2018; Onur et al., 2021), the *Drosophila* model system was leveraged (Freeman & Doherty, 2006; Ziegenfuss et al., 2012). An HD fragment *Drosophila* model where mHTT (*HTT^231NT128Q^*) is expressed in a pan-glia (*repo*) driver was crossed with lines that have a knockdown of the fly ortholog to *NFIA, NfI.* The astrogliogenesis transcriptional regulator, NfI, was evaluated due to its high orthology prediction (DIOPT) score (Y. Hu et al., 2011) with human NFIA and high activity in the larval ventral nerve chord, suggesting it may have an orthologous function in flies. Using a climbing assay (Al-Ramahi et al., 2018), mHTT flies exhibited a significant deficit in climbing compared to control flies (**Figure 8G**). The mHTT-induced climbing deficit was improved in lines with NfI knockdown in all cells (*NfI* KD) or in only glia (*NfI* glia KD) by 7 days and rescued to control levels by 10 days (**Figure 8G**). Therefore, one of the astrogliogenesis transcriptional regulators predicted from dysregulated HD mouse and human astrocyte transcriptomic analysis demonstrates that perturbation of astrogliogenesis can improve HD functional deficits *in vivo*.

## Discussion

Unbiased single cell approaches to define cell states across the landscape of thousands of individual cells—especially for cell types such as astrocytes whose function span a wide range of biological processes—provides a unique window into the cell states that arise during disease (Trapnell, 2015). The goal of our studies was to identify and compare astrocyte transcriptional cell states in specific systems and time points, their transitions in unaffected control and HD systems, and to elucidate the regulatory signaling involved. We investigated transcriptome signatures from human iPSC-derived astrocytes and astrocytes from a transgenic mouse model to elucidate common and species-specific cell states, as well as aberrant cell states that exist in both human and mouse HD astrocytes. Both models demonstrated potential loss of cell states involved in neural support, glutamate signaling, ion-binding, and neural development. Both also suggest astrocyte developmental alterations that are predicted to be regulated by astrogliogenesis transcription factors. HD human astrocytes exhibited a unique activation of ECM and cytoskeletal signaling that was not observed in the R6/2 astrocyte nuclei. While most of the transcriptional cell states we identified in both human and mouse data were represented in HD and control samples, with shifts in percent composition in each condition, the HD iAstro data revealed a complete loss of a unique control cell state that showed high expression of *NEAT1,* a gene associated with the stability of nuclear paraspeckles and thought to function through interactions with RNA binding and other proteins and RNAs, thus affecting transcription (Clemson et al., 2009; Hirose et al., 2014). *NEAT1* has also been implicated in striatal neurons from HD patients and mouse models, with its overexpression protecting from mHTT-induced cytotoxicity (Cheng et al., 2018). The R6/2 astrocytic dysregulation that overlapped between regions and with the human data showed an increase in severity between 8 and 12 weeks, suggesting that astrocyte dysregulation is progressive as described previously for bulk RNA-seq analysis of purified astrocytes (Diaz-Castro et al., 2019); however, there were unique differences found in 8-week cortical astrocytes involving SP1 signaling and lipid metabolism, which may represent unique age-dependent cortical changes or early changes that give rise to the dysregulation of molecular processes that were dysregulated at 12 weeks. Overall, the dysregulated astrocytic states identified here are anticipated to induce dysfunctional brain homeostasis and contribute to HD pathogenesis.

Single-nuclei technology has allowed us to identify diverse astrocyte states within a heterogeneous astrocyte population through unsupervised computational clustering that may not have been identified using traditional bulk transcriptomic analysis (X. Hu et al., 2016). Homeostatic astrocyte states identified from control iAstros and NT mouse astrocytes, compared to HD iAstros or R6/2 mouse astrocytes, included activation of signaling pathways previously suggested to play roles in astrocyte functions. Synaptogenesis signaling and CREB signaling were significantly activated in NT-enriched striatal and cortical astrocyte states along with calcium-binding activation in the cortex. In control human iAstros, several synaptogenesis-related genes were among the top upregulated cluster markers (*SPARCL1*) in control iAstro state cluster 0. Similarly, glutamate receptor signaling was significantly activated across an NT-enriched striatal mouse astrocyte state relative to R6/2 striatal astrocyte clusters. Overall, snRNA-seq of control iAstros and non-transgenic mouse astrocytes contained terms implicating synaptogenesis, glutamate signaling, and other neural supportive transcriptional cell states that are typical of homeostatic astrocytes.

### HD Human iAstro and R6/2 Astrocyte Phenotype Comparisons

The commonly dysregulated astrocyte state across HD human iAstros and R6/2 mouse astrocytes was the inhibition of glutamate signaling. This phenotype is consistent with previous studies suggesting that glutamate handling is a critical feature of excitotoxicity in HD (Choi, 1988; Cross et al., 1986; Greenamyre, 1986). Astrocytes terminate the action of neurotransmitters, like glutamate and gamma-aminobutyric acid (GABA), through uptake at the tripartite synapse. In the case of glutamate, astrocytes help clear this excitatory neurotransmitter from the synaptic cleft through transporters (GLT1 and GLAST) and with the help of transmembrane glutamate receptors (GluR1 and GluR2) that form ligand-gated ion channels (Lehre et al., 1995; Rothstein et al., 1994, 1996). Within the cytosol of astrocytes, glutamate is then degraded into the neuro-inactive state, glutamine, via glutamine synthetase (GLUL) in an ATP-dependent manner; each of these were downregulated in our present study. These changes were validated by decreased protein levels of the glutamate transporter GLAST, in human HD iAstros upon FACS isolation, and AMPA glutamate receptor GluR1 in GLAST-positive HD iAstros compared to control. Furthermore, these alterations lead to functional impairment though decreased glutamate receptor signaling, which correlated with decreased glutamate uptake in HD iAstros. This dysregulation is present in R6/2 and other models of HD and may involve potassium ion channel Kir4.1 in R6/2 mice (Bradford et al., 2009; Diaz-Castro et al., 2019; Faideau et al., 2010; Garcia et al., 2019; Jiang et al., 2016; Khakh et al., 2017; Lee et al., 2013). The evidence of dysfunctional glutamate regulation in our adult-onset HD iAstro lines is consistent with prior studies demonstrating a similar phenotype in juvenile-onset HD iPSC-derived astrocytes (Garcia et al., 2019), and with loss of GLT1 expression in mice with HTT-160Q selectively expressed only in astrocytes (Bradford et al., 2009). Our findings further support a cell autonomous component of aberrant glutamate signaling inhibition in astrocytes in HD with functional consequences in human cells.

Axonal guidance was an inhibited pathway predicted in one of the unique HD-enriched iAstro cell states which overlaps with our striatal R6/2 astrocytes. This neural developmental pathway is necessary to guide neurons to their correct targets for proper synaptogenesis, therefore inhibition of this critical process would alter neuronal circuitry homeostasis. Moreover, both HD iAstro-enriched cluster 3 and R6/2-enriched striatal cluster 4 appeared to be in a more immature, proliferative cell state. Aberrant activation of cell cycle-related signaling has been demonstrated from HD mice (Conforti et al., 2013; Luthi-Carter et al., 2002), HD iPSC-derived neural models (HD iPSC Consortium, 2017; Mattis et al., 2014; Ring et al., 2015; Smith-Geater et al., 2020), and HD patient tissue (Barnat et al., 2020; Hickman et al., 2021). Osipovitch et al also demonstrated a decrease in GFAP expression in HD patient ESC-derived astrocytes that reflects impaired astrocytic differentiation (Osipovitch et al., 2019).

Developmental alterations in HD iAstros and decreased mature astrocyte markers in a striatal R6/2 astrocyte subpopulation may be a common cell state associated with mHTT-induced decreased astrogliogenesis. Transcription factor analysis and protein validation in HD iAstros identified downregulation of early astrogliogenesis factors—SOX9 and ATF3 (Kang et al., 2012; Tiwari et al., 2018). These genes may be driving maturation impairments in HD astrocytes, and subsequently causing alterations in astrocyte function as these immature cell states also showed decreased expression of functional genes. HES1 overexpression was one of the most significantly enriched transcription factors in R6/2 12-week astrocytes along with the dysregulation of several HES genes in HD iAstros, consistent with epigenetic dysregulation of a HES family transcription factor (HES4) associated with striatal degeneration in HD post-mortem cortex (Bai et al., 2015). The observed molecular phenotypes may also be due to a loss of astrocyte identity rather than a developmental alteration, as suggested in previous studies (Langfelder et al., 2016); however, the human HD iAstros data suggest a developmental phenotype given the decreased propensity for HD iPSCs to differentiate into GLAST-positive, morphologically mature astrocytes, and the likelihood that this iPSC model recapitulates early developmental stages, similar to altered in vivo astroglial differentiation of human glial progenitor cells from HD embryonic stem cells (Osipovitch et al., 2019). Both hypotheses could be occurring as HD progresses and mature cells lose their identity while progenitor cells give rise to developmentally impaired astrocytes. Nonetheless, alterations in astrocyte functions are likely, in part, due to the lack of mature astrocyte characteristics. Rescuing these deficits may require the proper induction of transcriptional regulators that are involved in the development and maturation of astrocytes and the expression of mature functional genes.

NFIA is a master astrogliogenesis transcriptional regulator that acts as a co-regulator with SOX9 in early astrogliogenesis (Stevanovic et al., 2021) and ATF3 in later astrogliogenesis (Tiwari et al., 2018). We assessed perturbation of NFIA in HD animals. When the fly ortholog to NFIA, NfI, was knocked down in either all cells or in glia of a fly model of HD, the functional capacity for the mHTT repo flies to climb was significantly improved. An interpretation of this is that if a downregulated gene is knocked down and the phenotype is rescued, that gene may be downregulated as a compensatory mechanism (Onur et al., 2021). Another interpretation may be due to the differences in timepoints assessed across the three HD models utilized in this study. The autophagic balance of these transcription factors, like many others in HD, may be altered differently depending on disease stage due to mHTT-induced alterations in clearance and accumulation of such proteins (Hernandez et al., 2021; Jia et al., 2012; Ochaba et al., 2014). To this account, SOX9 was significantly decreased in HD iAstros, but only predicted to be a transcriptional regulator and not significantly decreased in R6/2 astrocytes. Investigations into additional timepoints across HD models are necessary for elucidation of this hypothesis. Overall, the astrogliogenesis transcriptional factors, like NFI, are dysregulated across multiple HD models and can be targeted for improved functional outcomes in a model of HD.

Astrocyte reactivity has been documented in HD mouse (Lin et al., 2001; Myers et al., 1991; Reddy et al., 1998; Yu et al., 2003) and human tissue (Sapp et al., 2001; Selkoe et al., 1982); however, the extent of cell-autonomous astrogliosis has not been extensively characterized. Surprisingly, control iAstros appeared to exhibit a more activated neuroinflammatory state compared to HD; however, this was a relatively small cell state in iAstro cluster 6, only encompassing 5% of the total nuclei sequenced. This suggests that control human iAstros may have a greater ability to mount normal inflammatory responses, including those that are a hallmark of *in vitro* cultured astrocytes (Haim et al., 2015). Neuroinflammatory signaling was enriched in R6/2 striatal and cortical genotype-specific differentially expressed genes, with minor activation in striatal 12-week astrocytes; however, looking closely into the cell states, NT-enriched cortical cluster 4 had a predicted activation of neuroinflammation signaling similar to iAstro control-enriched cluster 6. Consistent with this hypothesis, R6/2-enriched striatal cluster 4, which had the lowest expression of mature astrocyte markers, had a predicted inhibition of neuroinflammatory signaling, demonstrating a potential loss of cell function in HD. These data suggest that both activated and inhibited states may exist in HD and could be playing different roles in the disease. Furthermore, some of these neuroinflammatory states may only arise due to extrinsic signaling from other CNS cell types, as suggested by the lack of neuroinflammatory states in the HD iAstros. Since immune responses of astrocytes can induce detrimental and/or beneficial effects on neighboring cell types, it is unknown what consequences would result from loss of this cell state (Ding et al., 2021). Studies aimed at inducing reactivity through LPS or A1-activating factors (Liddelow et al., 2017), may help to determine if HD astrocytes are unable to mount appropriate responses. While a lack of inflammatory response in HD astrocytes contrasts with evidence in literature showing evidence of reactive astrocytes in HD mouse models (Lin et al., 2001; Reddy et al., 1998; Yu et al., 2003) and human post-mortem tissue (Faideau et al., 2010; Myers et al., 1991; Reddy et al., 1998; Sapp et al., 2001; Selkoe et al., 1982), more recent studies support a lack of traditional or A1-like reactive astrogliosis states in HD (Al-Dalahmah et al., 2020; Diaz-Castro et al., 2019; Jiang et al., 2016; Tong et al., 2014).

There were several dysregulated pathways common to HD human iAstros and R6/2 astrocytes; differences between systems may be due to multiple reasons including expression of an expanded repeat HTT exon 1 transgene in R6/2, while HD iAstros contain the full-length endogenous mHTT. Recent studies identified similarities and differences across astrocytes from truncated (R6/2) and full-length (zQ175) mHTT mouse and human stem cell derived-models (Benraiss et al., 2021) or post-mortem human striatal tissue (Diaz-Castro et al., 2019). Some similarities across mouse and human models in these studies and our study highlighted ion homeostasis, metabolism, neurotransmitter signaling (including glutamate receptor signaling), and astrocyte morphology (Benraiss et al., 2021; Diaz-Castro et al., 2019). Finally, species differences may also contribute as human astrocytes have greater development of astrocyte networks both in numbers and complexity, exhibiting significantly more multi-branched processes than rodent astrocytes (Khakh & Sofroniew, 2015; Oberheim et al., 2009). The increased ability of adult human astrocytes to respond to extracellular glutamate compared to mouse astrocytes is an example that suggests adult human astrocytes may have evolved an improved capability to respond to synaptic activity (Oberheim et al., 2009; Sun et al., 2013; Zhang et al., 2016).

### Unique HD Human iAstro Phenotypes

Signaling states unique to human HD iAstros included activation of actin cytoskeletal and integrin signaling. Additionally, HD iAstros had a decrease in matrix metalloproteinases (*MMP15 & 17*) that aid in ECM degradation. Altered actin cytoskeletal dynamics was validated by increased Phalloidin (F-actin) staining and the actin cross-linking protein, ACTN, in HD iAstros at the protein level. This gain-of-function cell state unique to human HD iAstros compared to R6/2 astrocytes, which suggests a cellular state transition that is most likely a cell-autonomous effect not influenced by the presence of other cell types that provide developmental cues for astrocytes. Altered actin cytoskeletal dynamics in HD have been implicated due to interaction with HTT, including the major actin monomer sequestering protein, profilin (Burnett et al., 2008; Goehler et al., 2004; Harjes & Wanker, 2003). The functional relevance of actin and integrin activation to disease pathogenesis has yet to be determined but may relate to altered morphology, adhesion, intracellular trafficking, and transcriptional regulation.

### Unique R6/2 Astrocyte Dysregulated Cell States

Cyclic adenosine monophosphate (cAMP)-mediated signaling was the most inhibited pathway in striatal and cortical R6/2 astrocytes at 12 weeks. This signaling pathway plays a vital role in cellular signaling, including the regulation of glucose and lipid metabolism. Interestingly, *Phkg1* was a downregulated gene common across both time points and both brain regions but was unique to R6/2. Since astrocytes are the primary storage site for glycogen in the brain (Sofroniew & Vinters, 2010), a decrease in *Phkg1* would reduce glycogen phosphorylase’s ability to break down glycogen in astrocytes needed for the high energy demand of neurons. Decreased glucose levels have been shown in HD astrocytes purified from mouse striatum (Polyzos et al., 2019) and decreased striatal metabolism in HD patients attributed to decreased glucose uptake through GLUT3 (Ciarmiello et al., 2006; McClory et al., 2014). In addition to disrupting neuronal activity, reduced glucose levels may impact downstream epigenetic and transcriptomic signatures.

We also described cellular states unique to the two brain regions assessed. Striatal R6/2 cell state differences highlighted changes in maturation and ion transport/binding astrocytic processes, while cortical R6/2 astrocytes mostly presented glycosylated cell state transition with an inhibited synaptogenesis signature. Striatal R6/2 astrocytes and HD iAstros clusters had a loss of astrocyte identity genes, while this molecular phenotype was less overt in cortical R6/2 astrocytes with fewer significant differentially expressed genes across genotype for 12-week and 8-week cortical astrocytes compared to striatal, which yielded less diversity in significant pathway dysregulation. This may correspond with more overt neurodegeneration in the striatum, compared to the cortex.

Taken together, our findings indicate that aberrant cell states exist in individual HD astrocytes that include inhibition of glutamate signaling and potential developmental alterations that may be regulated by mutant HTT’s interaction with astrogliogenesis transcription factors, ATF3, NFIA, and SOX9. In addition, loss of astrocyte maturation may contribute to inhibition of astrocytic functions, such as glutamate signaling, in human HD and R6/2 astrocytes. Knockdown of NFI in HD flies rescued motor deficits and may suggest that restored protein levels of this key astrogliogenesis transcription factor would be neuroprotective, possibly by allowing glial cells to reach full maturity if the protein is constitutively activated and functioning in the HD context. The identification of common and unique HD cell states across a heterogeneous population of human and mouse astrocytes provides directions for further mechanistic evaluation and greater information relating to potential therapeutic interventions targeting these dysregulated astrocytic states in HD as well as predicted outcomes across model systems.

## Acknowledgements

We thank HD patients and their families for their essential contributions to this research. We acknowledge the UCI Stem Cell Research Flow Cytometry Core and specifically, Vanessa Scarfone and Pauline Nguyen, for their FACS technical assistance. We thank the technical assistance of Jennifer Stocksdale for expansion of iPSC lines utilized in this paper and Marie Schulte-Bisping for assistance with Western blot optimization. We are grateful to Dr. Juan Botas at Baylor College of Medicine for generously providing HD fly lines for this manuscript. We also acknowledge Drs. Peter Donovan, Suzanne Sandmeyer, and Jeff Rothstein for helpful discussions. This work was supported by the following NIH grants: R35 NS116872 (LMT), R01834 NS089076 (LMT) and associated Research Supplement award (AMRO), P01 NS092525 (LMT), NRSA T32 Training Grant NS082174-05 (AMRO), 1RF1AG071683 (VS), and NS123207-01 (DVV). Additional support from NSF Bridge to the Doctorate Fellowship (AMRO), UCI Stanley Behrens Family Foundation (AMRO), UCI MIND startup funds (VS), CIRM Bridges to Stem Cell Program (CS), Hereditary Disease Foundation, Huntington’s Disease Society of America, and the Huntington’s Disease Community, Advocacy, Research & Education. This work was made possible in part, through access to the Genomics High Throughput Facility Shared Resource of the Cancer Center Support Grant (P30CA-062203) at the University of California Irvine and NIH shared instrumentation grants 1S10RR025496-01, 1S10OD010794-01, and 1S10OD021718-01.

## Author Contributions

AMRO, RGL, and LMT conceived and designed experiments. AMRO, KQW, and CS performed differentiations. AMRO, RM, and CS performed differentiation quality control analysis. WWP and EMA developed the differentiation paradigm. AMRO optimized the differentiation paradigm for selected cell lines. MSB performed the glutamate uptake assay and Sholl analysis. AL, JR, MSB, MN, EM, JR, and VS performed mouse experiments. AMRO and NGM performed Western blots. JW and RGL performed initial mouse transcriptomic data analysis. AMRO performed all mouse and human astrocyte transcriptomic analyses. SJH performed fly crosses and climbing assays in the laboratory of DVV. AMRO and MSB carried out data analysis. AMRO wrote the original draft of manuscript. AMRO, MSB, JR, WWP, RGL, and LMT edited the manuscript. LMT supervised the project and acquired funding.

## STAR Methods

### iPSCs

HD and unaffected control iPSCs were generated and characterized as described (HD iPSC Consortium, 2013, 2017; Lim et al., 2017; Mattis et al., 2013; Smith-Geater et al., 2020). All iPSC lines exhibited normal karyotypes by aCGH array performed by Cell Line Genetics (Madison, WI). The five iPSC lines were maintained on Matrigel hESC-qualified matrix (Corning) with mTeSR1 (STEM CELL Technologies). Once 60-80% confluent, iPSCs were passaged using a 5-10 min incubation with Versene (Gibco) at 37°C.

| Cell line | HTT CAG repeat | Sex | Reprogramming method | Fibroblast line | Clinical Notes |
| --- | --- | --- | --- | --- | --- |
| CS25iCTR18n6 (CTR18Q) | 18, 17 | Male | Non-integrating episomal | ND30625 | Clinically Normal; brother of affected sibling; mother affected and/or gene positive; Sibling(s) affected and/or gene positive; Niece is ND31038; HD family ND31846 |
| CS83iCTR33n1 (CTR33Q) | 33, 18 | Female | Non-integrating episomal | GM02183 | Clinically Normal; At risk (50%) for Huntington's Disease |
| CS04iHD46n10 (HD46Q) | 46, 26 | Male | Non-integrating episomal | UCI-HDF4 | Clinically Affected |
| CS03iHD53n3 (HD53Q) | 53, 18 | Male | Non-integrating episomal | UCI-HDF3 | Clinically Affected |
| CS81iHD71n3 (HD71Q) | 71, 17 | Female | Non-integrating episomal | GM04281 | Rigid form of HD, possible homozygote, onset at age 14 yrs; neurological exam 3/82 shows a hypokinetic variant of HD with dystonia and marked tremor, ataxic wide-based gait; maternal cousin to clinically normal GM04711 and clinically affected GM04777 |

### Astrocyte Differentiation

Five iPSC lines were differentiated into neural progenitor cells (NPCs) as described (Gibco #A1647801, MAN0008031). NPCs were cultured to 90-95% confluency, then seeded at 1e5/cm^2^ on hESC-qualified matrigel (Corning) for four passages before induction of astrocyte differentiation. All NPC lines exhibited normal karyotypes by aCGH array performed by Cell Line Genetics (Madison, WI). To begin astrocyte differentiation, passage four NPCs were seeded at 0.75e5/cm2 in Neural Expansion Medium (Gibco #A1647801, MAN0008031) and the following day (day 0), half the medium was replaced with astrocyte differentiation medium (ADM) 1. Half media changes were performed every other day for the remainder of the differentiation. All subsequent passages were performed by washing cells with HBSS without Mg Ca (Gibco) three times, then incubating in StemPro Accutase (Gibco) at 37°C for 10-15 min, counting using TC20 Automated Cell Counter (Bio-Rad) with trypan blue exclusion, and seeding on hESC-qualified matrigel (Corning) at 5e4/cm^2^ unless otherwise noted. On day 7, the cells were replated in ADM1. On day 15, the cells were replated in 50% ADM1, 50% ADM2. Subsequent half media changes were performed using ADM2. On day 30, day 34, and day 45, the cells were replated in 100% ADM3. On day 60, the cells were subject to astrocyte enrichment via FACS. QC immunostaining was performed at each passage time point for PSC, NPC, neural, and astrocytic markers.

| ADM1 Component | Company | Cat. # | Concentration |
| --- | --- | --- | --- |
| DMEM, high glucose | Life Technologies | 11995-065 | 100% |
| HI-FBS<sup>†</sup> | Life Technologies | 10082-147 | 1% |
| GlutaMAX [100X] | Life Technologies | 35050061 | 1X |
| N2 Supplement [100X] | Life Technologies | 17502048 | 1X |
| LIF* | STEMCELL Technologies | 78055 | 100ng/mL |
| CNTF* | Peprtech | 0117158 | 100ng/mL |
| bFGF* | STEMCELL Technologies | 78003.2 | 50ng/mL |

\*added to media fresh before media change
†Lot #: 1876835

| ADM2 Component | Company | Cat. # | Concentration |
| --- | --- | --- | --- |
| BrainPhys | STEMCELL Technologies | 05790 | 100% |
| SM1 Supplement [50X] | STEMCELL Technologies | 05711 | 1X |
| LIF* | STEMCELL Technologies | 78055 | 25ng/mL |
| CNTF* | Peprtech | 0117158 | 50ng/mL |
| bFGF* | STEMCELL Technologies | 78003.2 | 50ng/mL |

\*added to media fresh before media change

| ADM3 Component | Company | Cat. # | Concentration |
| --- | --- | --- | --- |
| DMEM, high glucose | Life Technologies | 11995-065 | 50% |
| Neurobasal | Life Technologies | 21103-049 | 50% |
| GlutaMAX [100X] | Life Technologies | 35050061 | 1X |
| Sodium Pyruvate [100mM] | Life Technologies | 11360070 | 1mM |
| SATO Supplement [100X] | - | - | 1X |
| N-Acetylcysteine* | Sigma-Aldrich | A9165 | 5µg/mL |
| huHBEGF* | R&D Systems | 259-HE | 5ng/mL |

\*added to media fresh before media change

### FACS Astrocyte-Enrichment

Day 60 cells were washed with HBSS without Mg Ca (Gibco) three times then lifted using a 10-15 min incubation of StemPro Accutase (Gibco) at 37°C. Cells were collected and strained through a 70um cell strainer then incubated with 100U/mL DNaseI (Qiagen) in FACS Buffer (0.5% BSA [Gibco] in PBS without Mg and Ca) for 10 min at room temperature. Cells were washed, collected, then stained for GLAST-APC (Miltenyi Biotec) in FACS Buffer for 15 min at 4°C in the dark. Cells were washed, 40um strained, then collected, then stained for DAPI in FACS Buffer for 5 min at room temperature in the dark. Cells were immediately sorted on the FACSAria Fusion Cell Sorter (BD Biosciences) into ADM3 with PenStrep (Gibco). GLAST-positive/DAPI-negative cells were replated at 0.38e6/well of a 6-well in ADM3 with 1x PenStrep. Following astrocyte-enrichment, the cells were cultured for another 7 days before harvest with half media changes every other day in ADM3 with 1x PenStrep followed by immunofluorescence quality control analysis for astrocyte markers.

### Immunofluorescence

Cells were fixed using 4% paraformaldehyde (Electron Microscopy Sciences) in PBS for 10 min at room temperature. Depending on antigen, cells were permeabilized with 0.3% Triton X-100 (Sigma-Aldrich) in PBS for 10 min at room temperature. Cells were blocked for 1 hr in 10% goat serum (Gibco), 1% BSA (Gibco), 0.1% Triton X-100 in PBS for 1 hr at room temperature. Cells were incubated with primary antibodies overnight at 4°C and then washed three times with PBS before incubation in 1:1000 Alexa-Fluor 488, 594, 647 secondary antibodies (Invitrogen) for 1 hr at room temperature and subsequently incubated with Hoechst 33342 (Sigma-Aldrich) for 10 min at room temperature. Stained slides were mounted with Fluoromount-G (Southern Biotech). Cells were imaged with a Nikon T-E fluorescence microscope at 10x and 20x. Analysis of images were performed by Cell Profiler (Carpenter et al., 2006).

| Primary Antibody | species | dilution | Company | Cat. # |
| --- | --- | --- | --- | --- |
| OCT4 | Mouse | 1:500 | Millipore | MAB4401 |
| SOX2 | Rabbit | 1:500 | Millipore | AB5603 |
| Nestin | Mouse | 1:1000 | Millipore | MAB5326 |
| PAX6 | Rabbit | 1:250 | Abcam | ab5790 |
| DCX | guinea pig | 1:500 | Millipore | AB2253 |
| TUJ1 | Mouse | 1:1000 | Millipore | MAB5564 |
| MAP2 | guinea pig | 1:1000 | Synaptic Systems | 188004 |
| GFAP | Chicken | 1:1000 | Abcam | ab4674 |
| ALDH1L1 | Rabbit | 1:400 | Abcam | ab87117 |
| S100β | Mouse | 1:500 | Abcam | ab4066 |
| GLAST | Mouse | 1:100 | Miltenyi Biotec | 130-095-822 |
| GLAST-APC | Mouse | 1:50 | Miltenyi Biotec | 130-123-555 |
| AQP4 | Rabbit | 1:100 | Abcam | ab81355 |
| GluR1 | Mouse | 1:1000 | Abcam | ab174785 |
| ATF3 | Rabbit | 1:1000 | Abcam | ab254268 |
| IgG | Mouse | As needed | Invitrogen | 02-6502 |
| IgG | Rabbit | As needed | Invitrogen | 02-6102 |

### Sholl and Morphometric Analysis

Individual iAstros (CTL and HD lines, n=25-40 cells, 5 differentiation replicates) were imaged on a Fluoview FV3000 Olympus Confocal Microscope at 40x magnification. iAstros were analyzed by Simple Neurite Tracer (SNT) in ImageJ (Arshadi et al., 2021), which provides 2D visualization of z-stack images. Images were z-projected to max intensity, changed to grayscale, and inverted for easier visualization. The SNT plug-in was accessed through Neuroanatomy, Deprecated, and then Simple Neurite Tracer. To build a primary path, a point was placed at the base of the path and then connected to the end of the path. Daughter paths were extended from these parent paths and to outline the full morphology. The primary and secondary number and path length were collected. Additionally, based on the paths, Sholl (Ferreira et al., 2014) was conducted to assess the complexity of each iAstros calculated by the number of intersections and concentric circles at 1um sequentially distant radii (Ferreira et al., 2014; Sholl, 1953).

### Glutamate Uptake Assay

Glutamate uptake from iAstrocytes was detected using an enzymatic, colorimetric glutamate assay kit (Abcam, ab83389). GLAST positive day 69 iAstrocytes cultured at 0.04e6 cells/cm^2^ were incubated with HBSS without Ca^2+^ and Mg^2+^ (without phenol red) for 30 min and then incubated with 200uM L-glutamic acid for 120 min. Cells were collected, counted, and processed according to manufacturer’s instructions.

### F-Actin In-Cell Western

Day 67 GLAST-positive astrocytes were seeded into a hESC-qualified matrigel coated 96-well plate at 0.04e6/cm^2^ for 3 technical replicates per biological (differentiation) replicate per cell line. Two days post seeding, cells were fixed using 4% paraformaldehyde (Electron Microscopy Sciences) in PBS for 20 min at room temperature. After washing cells three times with 0.1% Tween 20 (Sigma-Aldrich) in PBS, cells were permeabilized with 0.2% Triton X-100 (Sigma-Aldrich) in PBS for 30 min at room temperature. Cells were then blocked for 1 hr in 5% goat serum (Gibco), 3% BSA (Gibco) in PBS for 1 hr at room temperature. Cells were incubated with Phalloidin-iFluor790 CytoPainter (ab176763) at 1:1000 for 1 hr at room temperature, followed by washing three times with 0.1% Tween 20 (Sigma-Aldrich) in PBS. CellTag700 (LI-COR 926-41090) at 1:500 was added as a total protein stain (for normalization purposes) and incubated for 1 hr at room temperature. Cells were then washed three times with 0.1% Tween 20 (Sigma-Aldrich) in PBS, then solution was completely removed from wells. Wells were immediately imaged using the LI-COR Odyssey CLx and analyzed using LI-COR’s Empiria Studio Software.

| **Antibody** | **species** | **dilution** | **Company** | **Cat. #** |
| --- | --- | --- | --- | --- |
| GluR1 | Mouse | 1:1,000 | Abcam | ab174785 |
| ACTN | Mouse | 1:500 | Sigma-Aldrich | A5044 |
| ATF3 | Rabbit | 1:750 | Abcam | ab254268 |
| SOX9 | Rabbit | 1:500 | Sigma-Aldrich | AB5535 |
| IRDye® 700CW anti-Rabbit IgG | Goat | 1:20,000 | LI-COR Biosciences | 926-68071 |
| IRDye® 800CW anti-Mouse IgG | Goat | 1:20,000 | LI-COR Biosciences | 926-32210 |
| IRDye® 800CW anti-Rabbit IgG | Goat | 1:20,000 | LI-COR Biosciences | 926-32211 |

### Western Blots

Day 67 GLAST-positive astrocytes were broke in 150uL modified RIPA buffer, sonicated three times for 10 seconds at 40%, then spun down for 20 min at 4°C, 1000 rpm. Bradford (Thermo Scientific BioMate 3S UV-visible spectrophotometer) protein quantification was performed with cuvettes. Protein lysates were aliquoted at 5 ug for GluR1, ACTN, ATF3, and SOX9. Then equivalent amounts of loading buffer were added. Gels were ran at 150V using Bolt™ 4-12% Bis-Tris Plus Gels, 15-well gels with MOPS, transferred onto Nitrocellulose Membrane 0.45 µm at 10 V for 1 hr at room temperature. The membrane was rinsed in water and then let dry for 15 min. Membranes were rehydrated in water and then followed the Revert 700 Total Protein Stain protocol. Membranes were blocked in Intercept (TBS) Blocking Buffer for 1 hr at room temperature. Membranes were incubated in primary antibody (GluR1, ACTN, ATF3, and SOX9) and rotated at 4° overnight. Membranes were then washed three times with TBST, then incubated in secondary antibody: GluR1 into IRDye® 800CW Goat anti-Rabbit IgG, ACTN into IRDye® 800CW Goat anti-Mouse IgG, ATF3 into IRDye® 800CW Goat anti-Mouse IgG, and SOX9 in IRDye® 700CW Goat anti-Rabbit IgG for 1 hr at room temperature. Membranes were washed three times with TBST while gently shaking for 1 hr, then washed two times with TBS at room temperature. Membranes were imaged using the LI-COR Odyssey CLx and analyzed using LI-COR’s Empiria Studio Software.

### R6/2 Mouse Model

All experimental procedures were in accordance with the Guide for the Care and Use of Laboratory Animals of the NIH and animal protocols were approved by Institutional Animal Care and Use Committees at the University of California Irvine (UCI), an AAALAC accredited institution. R6/2 mice have been described elsewhere in detail (Mangiarini et al., 1996). For this study, five-week-old R6/2 and non-transgenic (NT) male mice were purchased from Jackson Laboratories and housed in groups of up to five animals/cage under a 12-hr light/dark cycle with ad libitum access to chow and water. Mice were aged to 8, 10, or 12 weeks then euthanized by pentobarbital overdose and perfused with 0.01 M PBS. Striatum and cerebral cortex were dissected out of each hemisphere and flash-frozen for snRNA-seq (8 and 12 weeks) or immunohistochemistry (10 weeks).

### snRNA-seq

*Mouse:* Single nuclei from three mice per genotype per timepoint were isolated from half hemisphere striatum or cortex in Nuclei EZ Lysis buffer (Sigma-Aldrich, NUC101-1KT) and incubated for 5 min. Samples were passed through a 70μm filter and incubated in additional lysis buffer for 5 min and centrifuged at 500 g for 5 min at 4°C before two washes in Nuclei Wash and Resuspension buffer (1xPBS, 1% BSA, 0.2U/μl RNase inhibitor). Nuclei were FACS sorted using DAPI to further isolate single nuclei and remove additional cellular debris. These nuclei were run on the 10x Genomics® Chromium Single cell 3’ gene expression v3 platform. Libraries were QCed and sequenced on the NovaSeq 6000.

*iAstros:* Day 67 GLAST-positive single-cell suspensions were strained using a 70 µm strainer, washed using cold 0.08% BSA (Gibco) in PBS, and counted using TC20 Automated Cell Counter (BioRad) with trypan blue exclusion as described in the 10x Genomics® Demonstrated Protocol: Single Cell Suspensions from Cultured Cell Lines for Single Cell RNA Sequencing (Document CG00054). To obtain a single-nuclei suspension, cells were lysed using Nuclei EZ Lysis Buffer (Sigma Aldrich, NUC101-1KT) for 5 min on ice. Lysed cells were centrifuged 500 rcf for 5 min at 4°C, then washed by twice resuspending in 1% BSA (Gibco), 0.2 U/µL of Human Placenta RNase Inhibitor (NEB), in PBS and centrifuged again. After two washes, the nuclei were strained using a 40µm cell strainer, counted using the TC20 Automated Cell counter (Bio-Rad), resuspended at 1000 nuclei/µL in 0.08% BSA (Gibco) in PBS, then proceeded immediately with the 10x Genomics® Chromium Single cell 3’ gene expression v3 platform. Libraries were QCed and sequenced on the NovaSeq 6000.

### Data Processing

*Mouse:* A total of 109,053 cells with 6.1 billion reads were sequenced for the 24 samples (on average 4,544 cells per sample with ∼55.6K reads per cell). Alignment was done using the CellRanger pipeline v3.1.0 (10x Genomics) to a custom pre-mRNA transcriptome built from refdata-cellranger-mm10-1.2.0 transcriptome using cellRanger mkref. To filter out low-quality nuclei and low-abundance genes, nuclei with less than 200 genes or more than 6000 genes and percent mitochondrial reads aligning to more than 2% of mitochondria genes were excluded from the downstream analyses. UMI counts were then normalized in Seurat 3.0 and top 2000 highly variable genes were identified using FindVariableFeatures function with variance stabilization transformation (VST). UMI Count matrices were generated from BAM files using default parameters of cellRanger count command. The gene barcode matrices for each sample were imported into R using the Read10X function in the Seurat R package (Stuart et al., 2019) (v3.1.5). Replicates were combined using cellRanger aggr. Dimension reduction was used to visualize and explore major features of the data using PCA and UMAP. Clustering was performed in Seurat v3.1 using PC1-20 and 0.5 resolution. The cluster with highest expression of astrocyte markers (Slc1a2, Slc1a3, Aldh1l1, S100b) were computationally subset to isolate a single astrocyte cluster for each dataset. Subclustering of astrocyte clusters was performed using PC1-20 and 0.5 resolution.

*iAstros*: A total of 37,170 cells with 2.3 billion reads were sequenced for the 4 samples (on average 9,293 cells per sample with ∼61K reads per cell). Fastq files were quality controlled and aligned to the GRCh38 reference transcriptome to obtain a gene count matrix using Cell Ranger pipeline v3.1.0 (10x Genomics). To filter out low-quality nuclei and low-abundance genes, nuclei with less than 200 genes or more than 6000 genes and percent mitochondrial reads aligning to more than 5% of mitochondria genes were excluded from the downstream analyses. UMI counts were then normalized in Seurat 3.1 and top 2000 highly variable genes were identified using FindVariableFeatures function with variance stabilization transformation (VST). After quality control and expression matrix formation, normalization was performed using UMI, and log-transformation was used to control variance. Seurat integration was used to combine datasets from patient-derived cell lines. Dimension reduction was used to visualize and explore major features of the data using PCA and UMAP. Clustering was performed in Seurat v3.1 using PC1-20 and 0.2 resolution.

### Differential Gene Expression and Pathway Analysis

To identify gene expression differences between clusters and/or samples, a non-parametric Wilcoxon rank sum test was used with Bonferroni correction and 1% FDR cutoff using Seurat v3.1. Cell types were assigned to clusters based on known cell-type markers. Data will be deposited in GEO. QIAGEN’s Ingenuity Pathway Analysis (http://www.qiagen.com/ingenuity) and Enrichr (Chen et al., 2013; Kuleshov et al., 2016) were used for analysis of significantly differentially expressed genes (FDR<0.1).

### Immunohistochemistry and Confocal Microscopy

Three mice per genotype at 10 weeks of age were euthanized by pentobarbital overdose and perfused with 0.01 M PBS. Whole brains were drop-fixed in 4% paraformaldehyde (Electron Microscopy Sciences) in PBS overnight at 4°C, then 30% sucrose for 48 hrs at 4°C. Half brain was embedded in Tissue-Tek O.C.T. Compound (Fisher Scientific) and sectioned coronally at a section thickness of 14μm on a cryostat at −20°C. Tissue was collected as free-floating sections in PBS with 0.05% sodium azide stored at 4°C. For staining, antigen retrieval with sodium citrate buffer (10mM Sodium Citrate, 0.05% Tween 20, pH 6.0) at 80°C for 30 min was performed with floating tissue sections. The sections were permeabilized for 15 min in 1X PBS, 0.25% TritonX-100, and 3% BSA (Gibco), and then blocked in 1X PBS, 0.1% TritonX-100, 10% Donkey Serum (Gibco) and 1% BSA. Sections were then immediately placed in a primary antibody diluted mix in 1X PBS, 5% Donkey Serum, and 0.1% TritonX-100 overnight at 4°C. Sections were washed three times with 1X PBS before incubation in 1:500 Alexa-Fluor 488, 594, 647 secondary antibodies (Invitrogen) for 1 hr at room temperature and subsequently incubated with Hoechst 33342 (Sigma-Aldrich) for 10 min at room temperature. Sections were mounted on microscope slides, coverslipped with Fluoromount-G (Southern Biotech), and visualized and captured using a Fluoview FV3000 Olympus Confocal Microscope at 60x magnification.

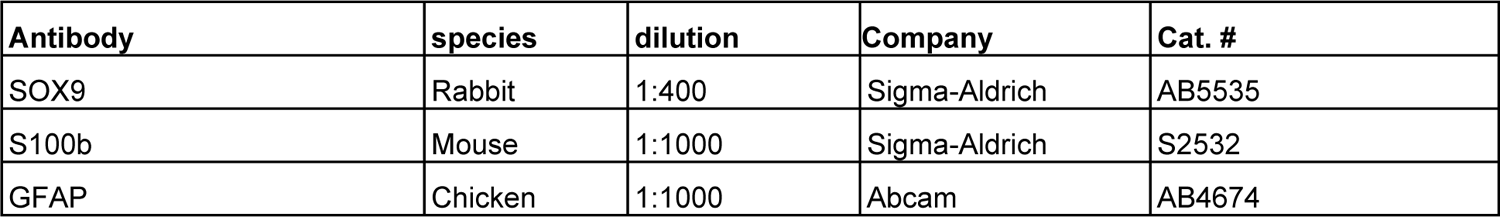

### Fly Husbandry

Fly lines were generated as described (Agrawal et al., 2005) or ordered from the Bloomington *Drosophila* Stock Center (BDSC) or Vienna *Drosophila* Stock Center (VDSC). A 231 amino acid N-terminal fragment of *mHTT* with 128 glutamines was expressed using a pan-glial (*repo*) driver. This line was generated in the Botas lab at Baylor College of Medicine and kindly provided as a gift. Flies were maintained at 18°C or 23°C on standard fly food (molasses, yeast extract, and agar).

| Gene | Genotype | Source | ID number |
| --- | --- | --- | --- |
| *HTT* repo | w; + ; HTT [231NT-128Q] - repo-gal4 / gal 80-TM6B | Botas lab |  |
| Control | Non-targeting hpRNA | VDSC | 13974 |
| *Nfl* KD | y[1]; Mi{y[+mDint2]=MIC}Nfl[MI06280] | BDSC | 60792 |
| *Nfl* glia KD | y[1] v[1]; P{y[+t7.7] v[+t1.8]=TRiP.JF02219}attP2 | BDSC | 31929 |

### Fly Climbing Assay

Virgin female flies harboring the *mHTT* transgene or control flies (VDSC 13974) were crossed to males harboring experimental alleles. Experimental crosses were raised and maintained at 25°C. Only female progeny were assessed using the climbing assay. Anesthetized using carbon dioxide, female progeny were sorted into vials used for experimentation without food in groups of 9 to 10 flies with 10 replicates (biological replicates). The percent of flies that climbed to or past 5cm in 10 seconds at room temperature were quantified. Tapping was used to elicit negative geotaxis to force flies to the bottom of the vials with 3 rapid succession taps, three times, over 2 seconds. Flies were allowed to climb for 10 seconds, and a picture was taken against a white background with a mark parallel to the lab bench at 5cm from the bottom of the vial. After a 1-minute recovery, the 10 second climbing trial was repeated for a total of five times (technical replicates). Flies were climbed at days 7 and 10 post-eclosion.

