## Supplemental Figures for "Integrated transcriptome analysis of Huntington’s disease iPSC-derived and mouse astrocytes implicates dysregulated synaptogenesis, actin, and astrocyte maturation"

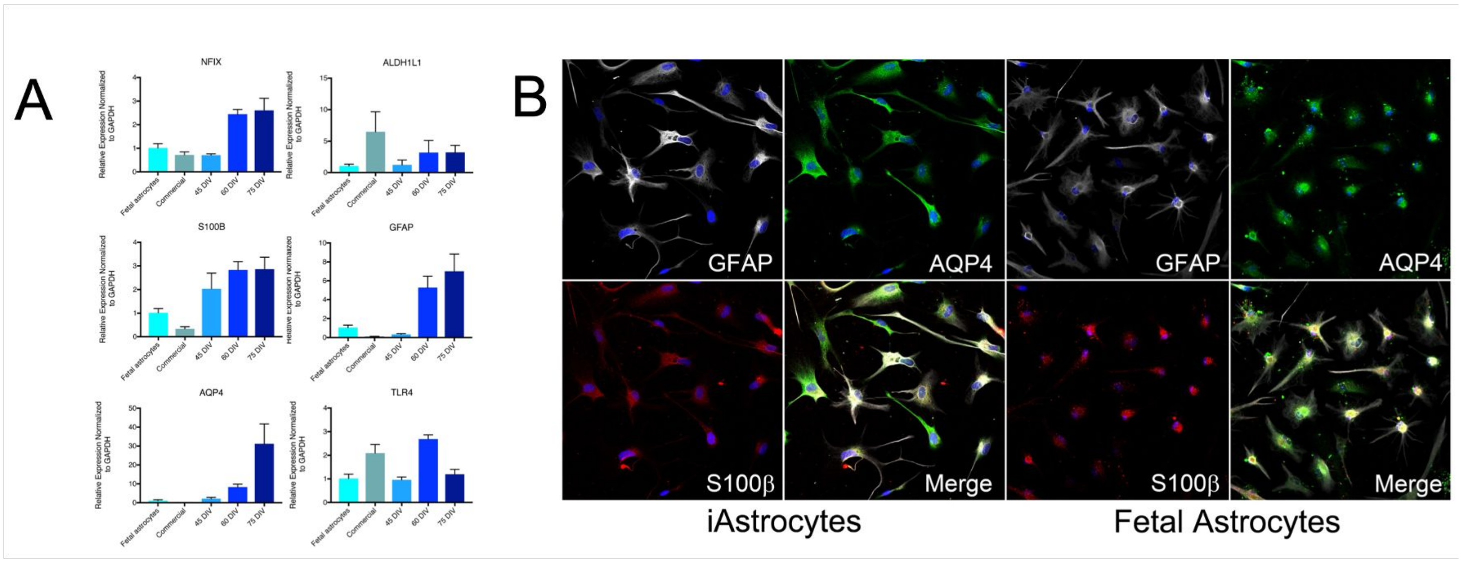


**Figure S1. iAstro differentiation compared to other human astrocyte cell lines.** (**A**) Quantitative PCR on astrocyte marker transcripts compared to human fetal and commercially available astrocytes from FUJIFILM Cellular Dynamics, Inc. (**B**) Immunocytochemistry of astrocyte marker proteins compared to human fetal astrocytes.

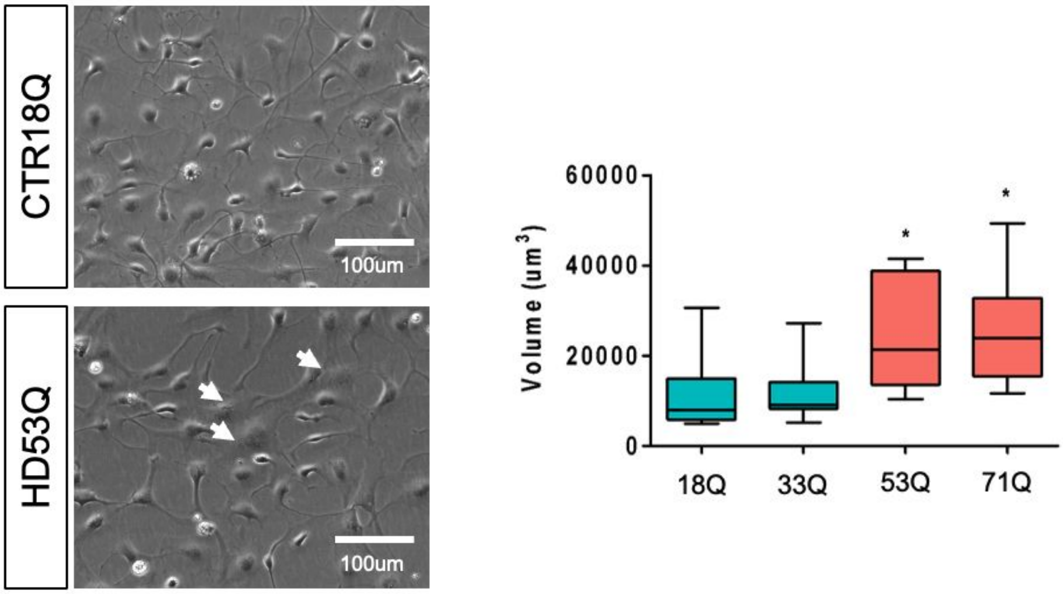


**Figure S2. HD and control iAstro morphology differences.** (**A**) Representative phase contrast images of unsorted day 60 iAstros morphology. HD iAstro white arrows highlight HD iAstros with enlarged cell bodies. (**B**) Quantification of cellular volume by confocal z-stacks of GFAP-positive iAstros at day 60. n = 3 biological (differentiation replicates) *p<0.05

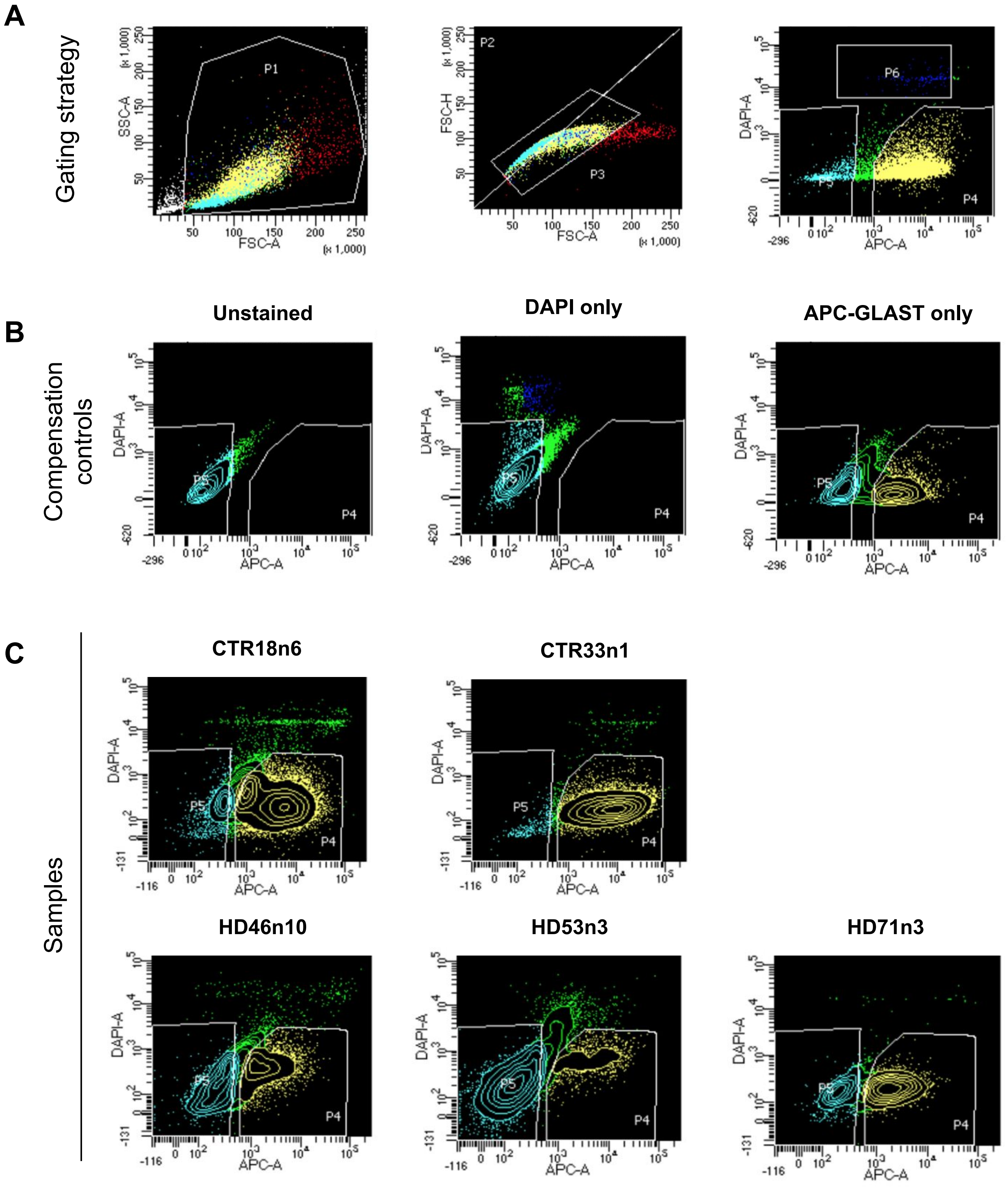


**Figure S3. Representative FACS Plots.** (**A**) Gating strategy to remove cellular debris and select single cells. (**B**) Compensation controls for unstained cells and single-stains of DAPI only and APC-GLAST only. (**C**) Representative FACS plot for each day 60 iAstro cell line. Population 4 (P4; yellow) was collected and used for subsequent assays as GLAST-positive iAstros.

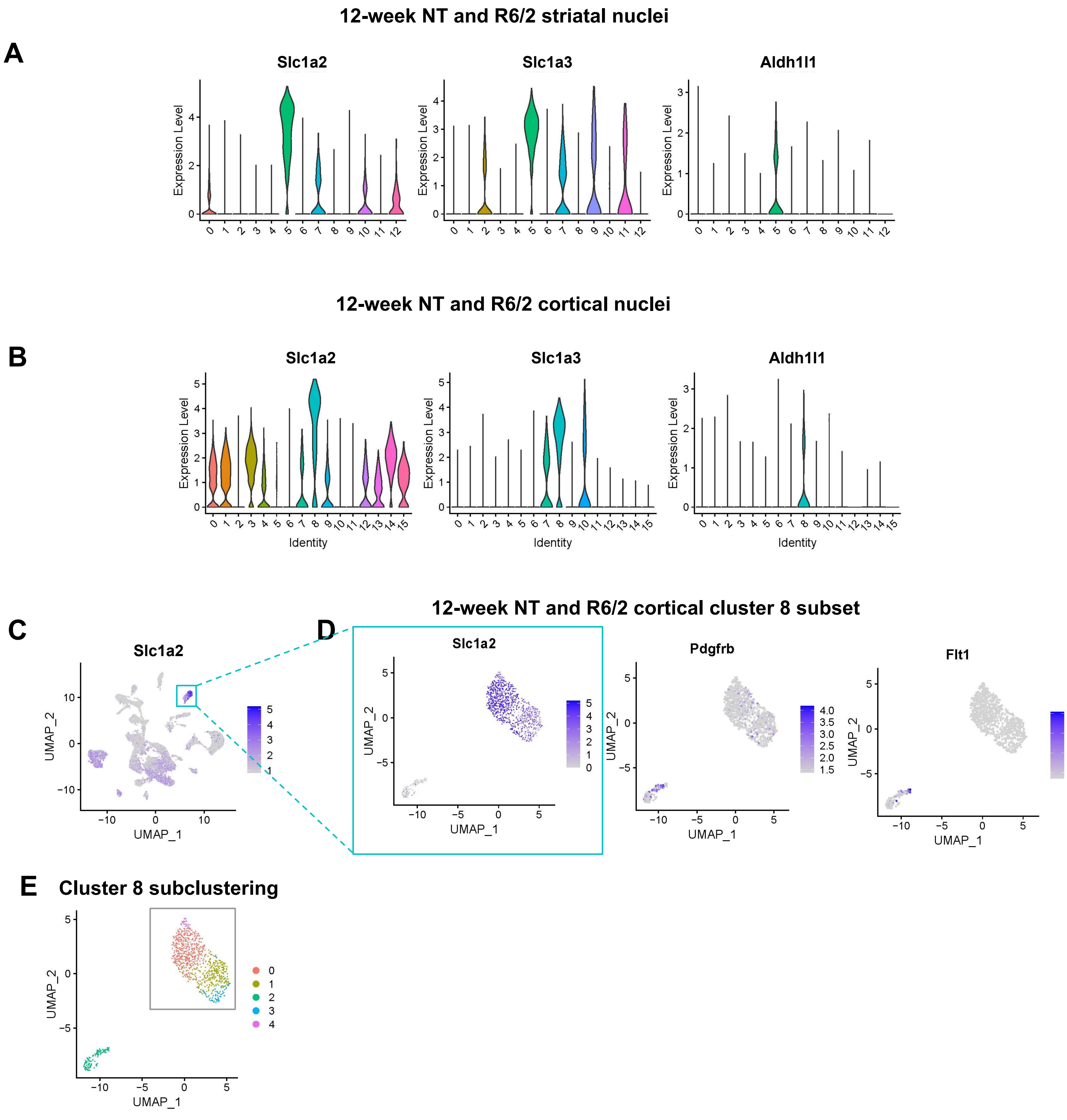


**Figure S4. Astrocyte cluster selection for NT and R6/2 12-week striatal and cortical snRNAseq.** (**A**) Violin plots of astrocyte markers depict striatal cluster 5 as the astrocyte cluster by highest astrocyte marker expression. (**B-D**) Violin plots (B) and UMAP visulization (C) of astrocyte markers depict cortical cluster 8 as the astrocyte cluster by highest astrocyte marker expression. (**D**) Subset of cortical cluster 8 with expression astrocyte markers and vascular markers shows multiple cell types. (**E**) Clustering analysis of cortical cluster 8 deomstrates subcluster 2 (green) contains significantly different gene expression compared to other subclusters. Subclusters 0, 1, 3, 4 (gray box) were subset and deemed the revised cortical astrocyte cluster for subsequent cortical analyses.

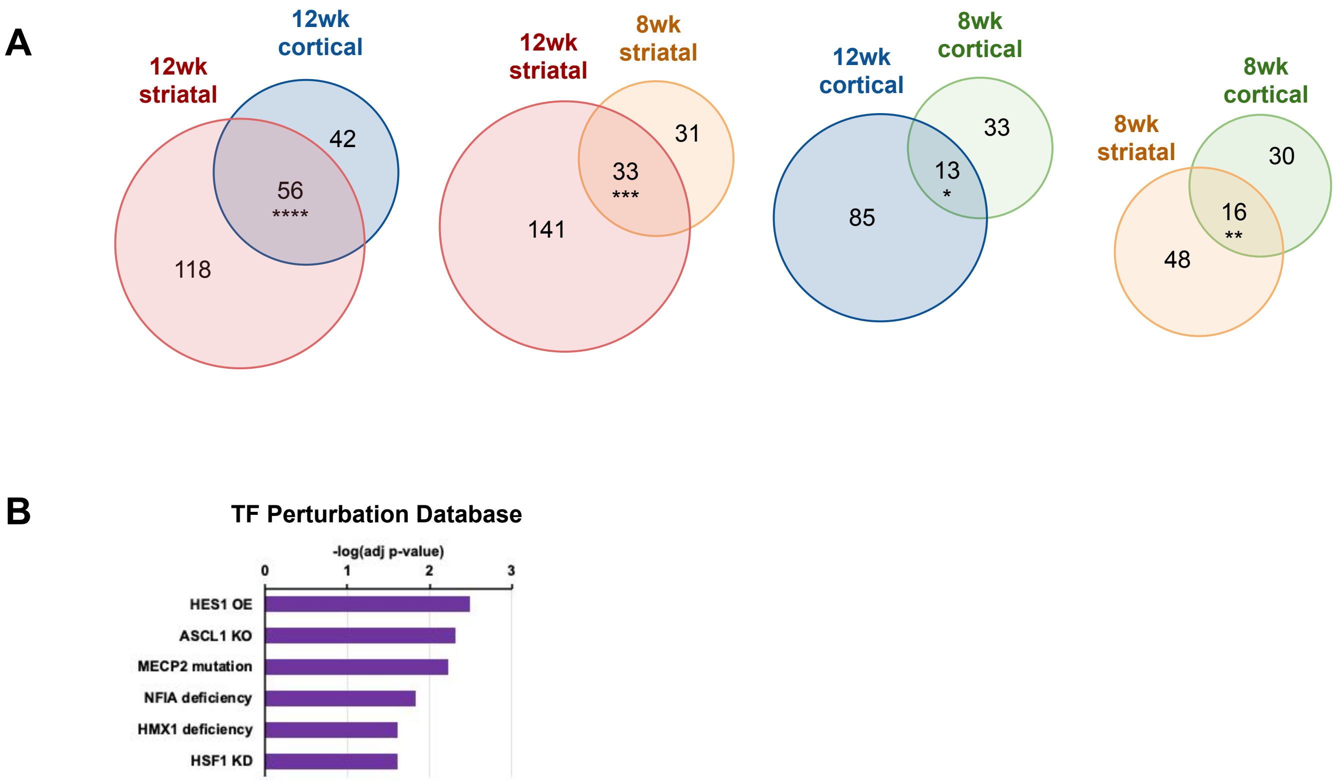


**Figure S5. R6/2 astrocyte differentially expressed gene overlaps and transcription factor enrichment.** (**A**) Venn diagrams of significant differentially expressed genes (DEGs) in R6/2 mouse astrocytes across time points and brain region (Exact hypergeometric probability calculated using 56 DEG overlap for 12 week striatum [174] vs 12 week cortex [98] ****p < 7.190e-113, 33 DEG overlap for 12 week striatum [174] vs 8 week striatum [64] ***p < 6.427e-64, 13 DEG overlap for 12 week cortex [98] vs 8 week cortex [46] *p < 7.335e-25, 16 DEG overlap for 8 week striatum [64] vs 8 week cortex [46] **p < 2.281e-35 using 46,206 as the number of genes with nucleotide sequence data in the mouse genome database). (**B**) Overlap of R6/2 12wk striatal and 12wk cortical astrocyte DEGs predicted transcription factor regulation using Enrichr.

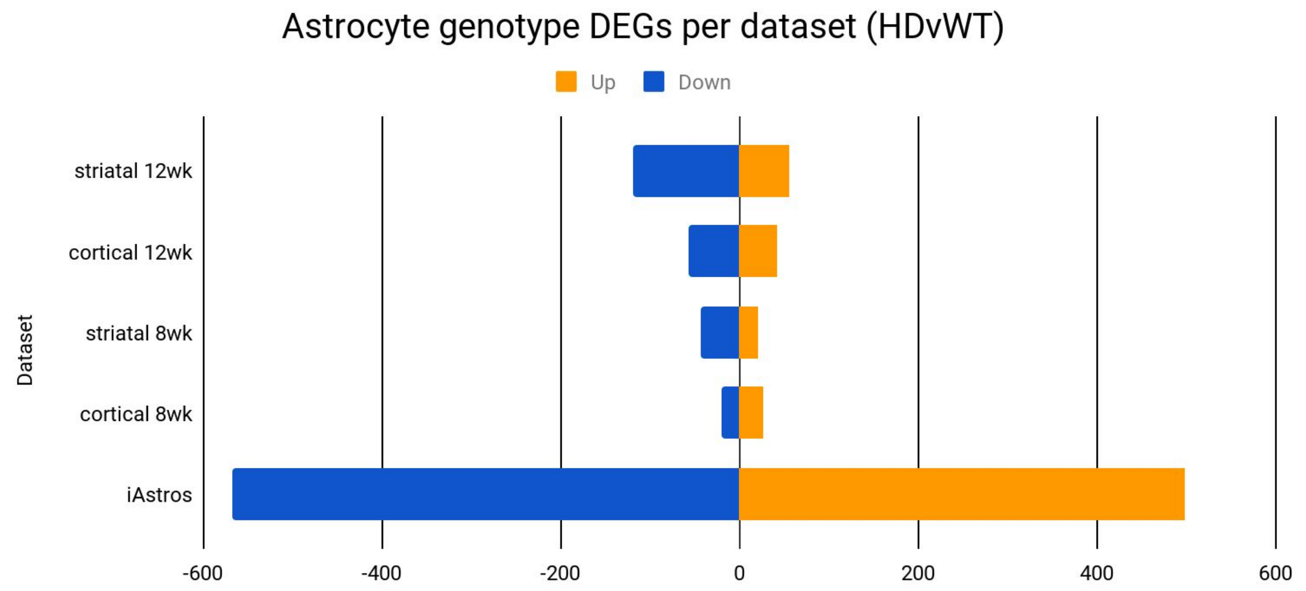


**Figure S6. Genotype DEG counts.** R6/2 astrocyte and HD iAstro significant differentially expressed gene count. Orange represents an upregulated gene and blue represents a downregulated gene count.
